# Sex differences in microRNA expression in first and third trimester human placenta

**DOI:** 10.1101/2021.05.13.444056

**Authors:** Amy E. Flowers, Tania L. Gonzalez, Nikhil V. Joshi, Laura E. Eisman, Ekaterina L. Clark, Rae A. Buttle, Erica Sauro, Rosemarie DiPentino, Yayu Lin, Di Wu, Yizhou Wang, Chintda Santiskulvong, Jie Tang, Bora Lee, Tianyanxin Sun, Jessica L. Chan, Erica T. Wang, Caroline Jefferies, Kate Lawrenson, Yazhen Zhu, Yalda Afshar, Hsian-Rong Tseng, John Williams, Margareta D. Pisarska

## Abstract

Maternal and fetal pregnancy outcomes related to placental function vary based on fetal sex, which may be the result of sexually dimorphic epigenetic regulation of RNA expression. We identified sexually dimorphic miRNA expression throughout gestation in human placentae. Next-generation sequencing was used to identify miRNA expression profiles in first and third trimester uncomplicated pregnancies using tissue obtained at chorionic villous sampling (n=113) and parturition (n=47). Sequencing and differential expression (DE) analysis identified 432 mature miRNAs expressed in the first trimester female, 425 in the first trimester male, 400 in the third trimester female, and 508 in the third trimester male placenta (baseMean >10). Of these, 11 sexually dimorphic (FDR<0.05, baseMean >10) miRNAs were identified in the first and 4 miRNAs were identified in the third trimester, including miR-361-5p, significant in both trimesters, all upregulated in females. Across gestation, 207 miRNAs were DE across gestation, common to both females and males, miR-4483, the most DE across gestation. There were twice as many female-specific differences across gestation as male-specific (44 miRNAs vs 21 miRNAs), indicating that miRNA abundance across human gestation is sexually dimorphic. Pathway enrichment analysis identified significant pathways that were differentially regulated in first and third trimester as well as across gestation. This work provides the normative sex dimorphic miRNA atlas in first and third trimester, as well as the sex independent and sex specific placenta miRNA atlas across gestation which may be used to identify biomarkers of placental function and direct functional studies investigating placental sex differences.

**Summary Sentence:** Sex dimorphism in miRNA expression is more pronounced in first compared to third trimester placenta, and there are twice as many female-specific gestational differences, indicating miRNA abundance across human gestation is also sexually dimorphic.

## Introduction

The effects of fetal sex on neonatal and pregnancy outcomes under varied conditions have been examined for decades [1-4]. Fetal growth is influenced by fetal sex, with males heavier at birth [4, 5]. Sexually dimorphic fetal outcomes more common in males than females include macrosomia, shoulder dystocia, cord dysfunctions, and low Apgar scores [4, 5]. Maternal outcomes more common with male fetuses include gestational diabetes, placental abruption, dysfunctional labor, prematurity, and assisted or cesarean deliveries [1, 4]. Some studies have shown an increased risk of preeclampsia and hyperemesis gravidarum with male fetuses [1, 6], although this has not been found consistently [7, 8]. The pathophysiologic mechanism driving sexually dimorphic outcomes remain poorly understood. As the primary route of communication between the fetomaternal unit, the placenta may drive many of these outcomes. The placental cellular structure, and thus genome derives from the fetus [9]. Therefore sex-specific outcomes may derive from sexually dimorphic gene expression in the placenta, which has been identified in the placenta [10, 11].

Epigenetic modifications, including post-transcriptional regulation, control overall gene expression, and phenotype. MicroRNAs (miRNAs) are small, single-stranded, noncoding RNA molecules, on average 22 nucleotides in length [12]. Of known miRNAs, approximately half originate from within introns of the protein coding genes they regulate, and can be co-transcribed [12]. miRNAs generally cause target messenger RNA to be degraded, or prevent translation to protein [13]; small changes in miRNA expression can result in multi-fold changes in gene expression[14]. In the placenta, miRNAs are important as early as trophectoderm development and implantation, some through export in exosomes [15-18], but these miRNA signatures change in the placenta throughout gestation during normal development [19-21]. In addition, there are two large miRNA clusters enriched in placenta, the chromosome 14 miRNA cluster (C14MC) and the chromosome 19 miRNA cluster (C19MC)[22, 23]. C14MC is a large, imprinted, maternally-expressed miRNA cluster, with several members predominantly expressed in placenta and epithelial tissues[24]. C19MC is a large, imprinted, paternally-expressed miRNA cluster whose members have highest expression in placenta and cancer, with relatively weak expression in other tissue[23-28]. In addition, two small clusters on chromosome 13 were recently identified to be uniquely present in the first trimester placenta [19, 29]. miRNAs have been identified in several pregnancy-related diseases, including pre-eclampsia [30-35], fetal growth [35-38], and gestational diabetes [39, 40], all sexually dimorphic pregnancy complications. However, sex differences and sex specific miRNA signatures in normal healthy gestations have not been defined.

Therefore, to identify potential biomarkers of placental function, it is imperative to first determine if sex differences in miRNA signatures exist in the first and third trimester placenta of normal healthy gestations. Furthermore, it is critical to identify miRNA signatures that are sex independent and sex specific to develop normative miRNA signatures. Therefore, we performed next-generation sequencing (NGS) and expression analysis to identify and compare sexually dimorphic miRNA expression in first and third trimester placentae of healthy pregnancies resulting in delivery. Furthermore, we identified miRNA signatures of common and sexually dimorphic miRNA expression across gestation for future development of biomarkers of placental disease early in gestation that are sex independent and also sex specific.

## Materials & Methods

### Study Population

The study population consisted of 157 singleton pregnancies between 2009 and 2018, including 113 with leftover chorionic villus sampling (CVS) tissue (58 female and 55 male), 47 with third trimester placenta collected at delivery (19 female and 28 male), and 3 patients with matched first and third trimester samples. All subjects were enrolled with informed written consent under IRB approved protocols (Pro00006806, Pro00008600). The cohorts of male and female fetuses were matched for parental age, maternal medical conditions, and fetal race and ethnicity at the time of enrollment, to minimize confounding factors that could impact outcomes. All pregnancies had a normal karyotype and resulted in the delivery of a viable infant.

### Collection of placental samples

Samples from the first trimester of pregnancy were collected at 10.0-14.5 weeks gestation during CVS procedures done for prenatal diagnosis. Samples used for research consisted of tissue which is normally discarded once enough tissue is obtained for prenatal diagnosis. Fetal-derived chorionic villi were cleaned and separated from any maternally-derived decidua. Samples from the third trimester of pregnancy were collected between 36.3-41.4 weeks gestation, after delivery of a viable neonate. Samples used for research consisted of tissue which would have otherwise been discarded. One centimeter cubed placental tissue samples were obtained immediately after delivery from the fetal side of the placenta near the site of cord insertion beneath the amnion, then separated into smaller pieces. Tissue samples were kept on ice and submerged in RNA*later* RNA stabilization reagent (QIAGEN, Hilden, Germany) within 30 minutes of collection, and stored in −80°C.

### Analysis of demographic data

For demographic analyses, 3 patients with matched first and third trimester samples were omitted, resulting in N=110 (57 female and 53 male) first trimester and N=44 (18 female and 26 male) third trimester samples for this analysis. Demographic data were collected including parental ages, races, and ethnicities, maternal pre-pregnancy body mass index, fetal sex, maternal medical history and medication use, pregnancy complications, mode of delivery, gestational age at delivery, and birth weight. Means and standard deviations were reported for continuous variables. T-test was used for normally distributed continuous variables, and the Wilcoxon rank-sum test for non-parametric data. Fisher’s exact test was also used as appropriate. Chi-square test was used for comparison of categorical variables.

### RNA extraction from chorionic villi

RNA extraction was performed from the first trimester placental samples utilizing a method optimized for extracting DNA and total RNA including small RNAs [10]. Briefly, tissue samples were thawed on ice with 600 µl of RLT Plus lysis buffer (QIAGEN) and 1% β-mercaptoethanol added to each sample, then homogenized by passing the tissue through progressively thinner needles (22G, 25G, and 27G) attached to an RNase-free syringe. Homogenates were loaded onto AllPrep spin columns and the remainder of sample processing was performed following manufacturer instructions using the AllPrep DNA/RNA/miRNA Universal Kit (QIAGEN). RNA was eluted with 30-45 µl of RNase-free water at room temperature and the elution was passed through the column twice to improve yields, as previously described [10, 41]. Equal numbers of male and female samples were processed each time to reduce batch effects. The average RNA integrity number (RIN) for sequenced samples was 8.87.

### RNA extraction from third trimester placenta

RNA extraction was performed from the third trimester placenta samples utilizing a method similar to that used for the RNA extraction from chorionic villi. However, to homogenize third trimester placental tissue, the tissue was sonicated in ice-cold RLT + 1% beta-mercaptoethanol buffer, using 5 second pulses on a low setting (#2) until tissue fragments were small enough to complete homogenization with RNase-free needles. The average RIN for sequenced samples was 8.84.

### Library preparation and miRNA sequencing

A miRNA sequencing library was prepared using the QIASeq miRNA Library Kit (QIAGEN, Hilden, Germany) from total RNA. A pre-adenylated DNA adapter was ligated to the 3’ ends of miRNAs, followed by ligation of an RNA adapter to the 5’ end. A reverse-transcription primer containing an integrated Unique Molecular Index (UMI) was used to convert the 3’/5’ ligated miRNAs into cDNA. After cDNA cleanup, indexed sequencing libraries were generated via sample indexing during library amplification, followed by library cleanup. Libraries were sequenced on a NextSeq 500 (Illumina, San Diego, CA) with a 1×75 bp read length and an average sequencing depth of 10.64 million reads per sample.

### Differential expression analysis of miRNAs

The demultiplexed raw reads were uploaded to GeneGlobe Data Analysis Center (QIAGEN) at https://www.qiagen.com/us/resources/geneglobe/ for quality control, alignment and expression quantification. Briefly, 3’ adapter and low quality bases were trimmed off from reads first using cutadapt v1.13 [42] with default settings, then reads with less than 16 bp insert sequences or with less than 10 bp UMI sequences were discarded. The remaining reads were collapsed to UMI counts and aligned sequentially to miRBase v21 mature, hairpin and piRNA (piRNABank) databases using *Bowtie* v1.2 [43, 44]. The UMI counts of each miRNA category were quantified, and then normalized by a size factor-based method in the R package *DESeq2* version 1.22.2 (Bioconductor) [45]. Data were averaged across all samples in each group for each respective miRNA (male and female) and were reported as baseMean. Eight miRNAs with no tissue expression were excluded from analysis (baseMean=0, miR-4776-3p and seven piRNAs). Next, the R package *FactoMineR* release 1.41 was used to conduct principal components analysis (PCA), which was used to investigate clustering and potential outliers. Each miRNA was fitted into a negative binomial generalized linear model, and the Wald test was applied to assess the differential expressions between two sample groups. Benjamini and Hochberg procedure was applied to adjust for multiple hypothesis testing, and significantly differentially expressed miRNA candidates were selected as those with a false discovery rate (FDR) less than 0.05. The chromosomal location of differentially expressed miRNAs was ascertained using miRBase v21 and *biomaRt* v2.45.8 R package with Ensembl release 91 [46, 47], and plotted against the number of miRNAs localized to the respective chromosome. For miRNAs derived from more than one chromosome, each miRNA precursor was counted separately in order to capture all chromosome sources (e.g. in barplots, miR-514a-3p is encoded by three precursors on chromosome X, and thus was counted three times). To avoid variable copy number regions, precursor duplications on the same chromosome were not included. For scatter plots, each miRNA was plotted only once per chromosome. The R package *gplots* v3.0.1.1 was used to generate a heatmap showing normalized expression for the 29 differentially expressed miRNAs (FDR<0.05). Finally, the R package *ggplot2* v3.1.1 was used to generate a factor analysis (FA) plot, plotting log transformed mean expression against fold change, and a volcano plot, plotting log transformed fold change against FDR.

### Predictive analysis of RNA targets for miRNAs with highest expression

Ingenuity Pathways Analysis (IPA) software’s microRNA Target Filter application (QIAGEN, Redwood City, CA, USA, http://www.qiagenbioinformatics.com/IPA) was used to generate a list of predicted target mRNAs based on sequence and experimental confirmation. Highly expressed miRNAs were included as inputs if they had baseMean>2000, and each sex and trimester was analyzed individually. RNA targets were only included if biochemically confirmed using human tissue or non-species specific methods (sourced from QIAGEN’s curated Ingenuity Knowledge Base[48], or the publicly available miRecords[49] or TarBase[50]), based on the TargetScan algorithm previously described[51]. After the list of target mRNAs was finalized, IPA’s Core Analysis function was used to test the hypothesis that the listed genes were part of canonical biological pathways, as previously described [10, 52, 53].

### Predictive analysis of RNA targets for sex different miRNAs

The procedure described above was applied for between-sex differentially expressed miRNAs (FDR<0.05, baseMean>10 in each sex), within-sex differentially expressed miRNAs (FDR<0.05, baseMean>10 in each trimester), and within-sex cluster-specific differentially expressed miRNAs (FDR<0.05, baseMean>10 in each trimester), with each sex and trimester analyzed separately.

### Heatmaps

Heatmaps with dendrograms of miRNAs versus samples were generated with matrices of log2(baseMean) data scaled and centered by rows. The plots were then generated with hierarchical clustering using the R package *gplots* v3.1.1. Heatmaps of gene enrichment data were generated using the R package *pheatmap* v1.0.12 with matrices of -log_10_(P) output from IPA Core Enrichment Analysis. For comparisons of -log_10_(P) values across rows in gene enrichment heatmaps, a difference of 4 or greater was considered the threshold at which visually distinct results were achieved.

## Results

### Cohort demographics and birth outcomes

Female and male singleton pregnancies in the first (N=113, 58 female and 55 male) and third trimester (N=47, 19 female and 28 male fetuses) were matched for maternal age, race, ethnicity, and pre-existing medical conditions. Principal component analysis (PCA) of placenta miRNA expression shows no distinct segregation by sex, although PCA by trimester shows a distinct segregation by first and third trimester placenta divided by principal component 1 (Supplemental File 1). Parental and fetal race and ethnicity were not significantly different between sexes in either first or third trimester (Table 1A). Maternal pre-pregnancy BMI was not significantly different between the sexes in either trimester. All pregnancies resulted in live births and there were no significantly different pregnancy complications between the sexes in either trimester cohort. Males were larger than females at birth in both sequencing cohorts, though only significant in the first trimester cohort.

**Table 1:** Demographics of study patient cohorts. Values shown as mean (standard deviation) or n (%) as applicable. Asterisk: denotes significance at p<0.05.

A) Comparison of Male and Female Fetuses in each gestation
|  | First Trimester (N=110) | Third Trimester (N=44) |
| --- | --- | --- |

|  | Female | Male | P | Female | Male | P |
| --- | --- | --- | --- | --- | --- | --- |
| **N** | 57 | 53 |  | 18 | 26 |  |
| **Maternal age, years** | 37.8 (2.9) | 37.5 (3.1) | 0.61 | 37.2 (2.5) | 37.3 (3.4) | 0.45 |
| **Paternal age, years** | 39.5 (4.6) | 39.4 (5.1) | 0.96 | 39.4 (4.8) | 38.0 (4.8) | 0.51 |
| **Maternal race/ethnicity** |  |  |  |  |  |  |
| Caucasian | 53 (93.0%) | 53 (100%) | 0.12 | 13 (72.2%) | 22 (84.6%) | 0.45 |
| Non-Hispanic | 55 (96.5%) | 52 (98.1%) | 1 | 14 (77.8%) | 21 (80.8%) | 1 |
| **Paternal race/ethnicity** |  |  |  |  |  |  |
| Caucasian | 54 (94.7%) | 50 (94.3%) | 1 | 15 (83.3%) | 22 (84.6%) | 1 |
| Non-Hispanic | 56 (98.3%) | 51 (96.2%) | 0.61 | 15 (83.3%) | 21 (80.8%) | 1 |
| **Fetal race/ethnicity** |  |  |  |  |  |  |
| Caucasian | 53 (93.0%) | 50 (94.3%) | 1 | 12 (66.7%) | 20 (76.9%) | 0.51 |
| Non-Hispanic | 55 (96.5%) | 51 (96.2%) | 1 | 13 (72.2%) | 20 (76.9%) | 0.74 |
| **Maternal pre-pregnancy BMI, kg/m2** | 22.0 (3.4) | 21.8 (3.4) | 0.54 | 23.9 (4.6) | 24.0 (4.8) | 0.98 |
| **Maternal preexisting medical conditions** |  |  |  |  |  |  |
| Hypertension | 0 (0%) | 0 (0%) | - | 0 (0%) | 0 (0%) | - |
| Diabetes | 0 (0%) | 0 (0%) | - | 0 (0%) | 0 (0%) | - |
| **Pregnancy complications** |  |  |  |  |  |  |
| Hypertension | 0 (0%) | 0 (0%) | - | 3 (16.7%) | 5 (19.2%) | 1 |
| Diabetes | 1 (1.8%) | 2 (3.8%) | 0.42 | 0 (0%) | 1 (3.9%) | 1 |
| Placenta previa | 0 (0%) | 0 (0%) | - | 0 (0%) | 1 (3.9%) | 1 |
| Placental abruption | 0 (0%) | 0 (0%) | - | 0 (0%) | 0 (0%) | - |
| **Hypertension management in pregnancy** |  |  |  |  |  |  |
| Anti-Hypertensives | 0 (0%) | 0 (0%) | - | 1 (5.6%) | 2 (7.7%) | 1 |
| Any Magnesium use (Ante- or Postpartum) | 0 (0%) | 0 (0%) | - | 2 (11.1%) | 4 (15.4%) | 1 |
| **Mode of delivery- Cesarean section** | 19 (33.3%) | 14 (26.4%) | 0.45 (chi2) | 7 (38.9%) | 8 (30.8%) | 0.59 (chi2) |
| **Gestational age at delivery, days** | 276.2 (7.0) | 276.3 (7.0) | 0.95 | 278.8 (5.2) | 275.0 (9.3) | 0.13 |
| **Birthweight, g** | 3342.0 (430.1) | 3536.0 (481.3) | 0.03* (ttest) | 3397.6 (409.5) | 3525.5 (504.1) | 0.48 |

B) Comparison of Sex Specific First and Third Trimester Samples
|  | **Female (N=75)** |  |  | **Male (N=79)** |  |  |
| --- | --- | --- | --- | --- | --- | --- |
|  | First | Third | P Val | First | Third | P Val |
| **N** | 57 | 18 |  | 53 | 26 |  |
| **Maternal age, years** | 37.8 (2.9) | 37.2 (2.5) | 0.34 | 37.5 (3.1) | 37.3 (3.4) | 0.87 |
| **Paternal age, years** | 39.5 (4.6) | 39.4 (4.8) | 0.79 | 39.4 (5.1) | 38.0 (4.8) | 0.27 |
| **Maternal race/ethnicity** |  |  |  |  |  |  |
| Caucasian | 53 (93.0%) | 13 (72.2%) | 0.03* | 53 (100%) | 22 (84.6%) | 0.01* |
| Non-Hispanic | 55 (96.5%) | 14 (77.8%) | 0.03* | 52 (98.1%) | 21 (80.8%) | 0.01* |
| **Paternal race/ethnicity** |  |  |  |  |  |  |
| Caucasian | 54 (94.7%) | 15 (83.3%) | 0.15 | 50 (94.3%) | 22 (84.6%) | 0.21 |
| Non-Hispanic | 56 (98.3%) | 15 (83.3%) | 0.04* | 51 (96.2%) | 21 (80.8%) | 0.04* |
| **Fetal race/ethnicity** |  |  |  |  |  |  |
| Caucasian | 53 (93.0%) | 12 (66.7%) | 0.01* | 50 (94.3%) | 20 (76.9%) | 0.053 |
| Non-Hispanic | 55 (96.5%) | 13 (72.2%) | 0.007* | 2 (3.8%) | 6 (23.1%) | 0.014* |
| **Maternal pre-pregnancy BMI, kg/m<sup>2</sup>** | 22.0 (3.4) | 23.9 (4.6) | 0.11 | 21.8 (3.4) | 24.0 (4.8) | 0.03* |
| **Maternal preexisting medical conditions** |  |  |  |  |  |  |
| Hypertension | 0 (0%) | 0 (0%) | - | 0 (0%) | 0 (0%) | - |
| Diabetes | 0 (0%) | 0 (0%) | - | 0 (0%) | 0 (0%) | - |
| Thyroid disorder | 1 (1.8%) | 2 (11.1%) | 0.14 | 2 (3.8%) | 4 (15.4%) | 0.087 |
| **Pregnancy complications** |  |  |  |  |  |  |
| Hypertension | 0 (0%) | 3 (16.7%) | 0.012* | 0 (0%) | 5 (19.2%) | 0.003* |
| Diabetes | 1 (1.8%) | 0 (0%) | 1 | 2 (3.8%) | 1 (3.9%) | 1 |
| Placenta previa | 0 (0%) | 0 (0%) | - | 0 (0%) | 1 (3.9%) | 0.55 |
| Placental abruption | 0 (0%) | 0 (0%) | - | 0 (0%) | 0 (0%) | - |
| **Hypertension management in pregnancy** |  |  |  |  |  |  |
| Anti-Hypertensives | 0 (0%) | 1 (5.6%) | 0.24 | 0 (0%) | 2 (7.7%) | 0.11 |
| Any Magnesium use (Ante- or Postpartum) | 0 (0%) | 2 (11.1%) | 0.055 | 0 (0%) | 4 (15.4%) | 0.003* |
| **Mode of delivery- Cesarean section** | 19 (33.3%) | 7 (38.9%) | 0.67 (chi2) | 14 (26.4%) | 8 (30.8%) | 0.72 (chi2) |
| **Gestational age at delivery, days** | 276.2 (7.0) | 278.8 (5.2) | 0.099 | 276.3 (7.0) | 275.0 (9.3) | 0.84 |
| **Birthweight, g** | 3342.0 (430.1) | 3397.6 (409.5) | 0.77 | 3536.0 (481.3) | 3525.5 (504.1) | 0.93 |

When comparing sexes across gestation, there were no significant differences in parental ages but there were significant differences in parental and fetal race and ethnicity (Table 1B). Pre-pregnancy BMI was greater in mothers of males in the third trimester cohort compared to the first trimester cohort. Maternal pre-existing medical conditions were not significantly different among the groups but there were 3 mothers who developed hypertension in the third trimester female cohort (some of which required magnesium) and 5 mothers in the third trimester male cohort, compared to none in the first trimester cohorts, which was significantly different.

### MiRNAs expressed by male and female placentae throughout gestation

In first trimester, female placenta expressed 432 mature miRNAs derived from 484 precursors (baseMean >10), and male placenta expressed 425 mature miRNAs derived from 467 precursors (baseMean>10), from all 22 autosomes and the X chromosome (Figure 1Ai). No annotated miRNAs from the Y chromosome were identified in miRBase release 21. The majority of expressed miRNAs in both sexes originate with similar distribution from chromosome 19 (Female, 123/484 [25.41%]; Male, 123/467 [26.34%]), chromosome 14 (Female, 103/484 [21.28%]; Male, 104/467 [21.28%]), the X chromosome (Female, 93/484 [19.21%]; Male, 91/467 [19.49%]), and chromosome 1 (Female, 67/484 [13.84%]; Male, 64/467 [13.70%]). Among the most highly expressed miRNAs (baseMean >2000), 80 miRNAs were expressed in females and 78 in males, predominantly from chromosome 19 (Female, 61/80 [76.25%]; Male, 61/78 [78.21%]; Figure 1Aii).

**Figure 1:**
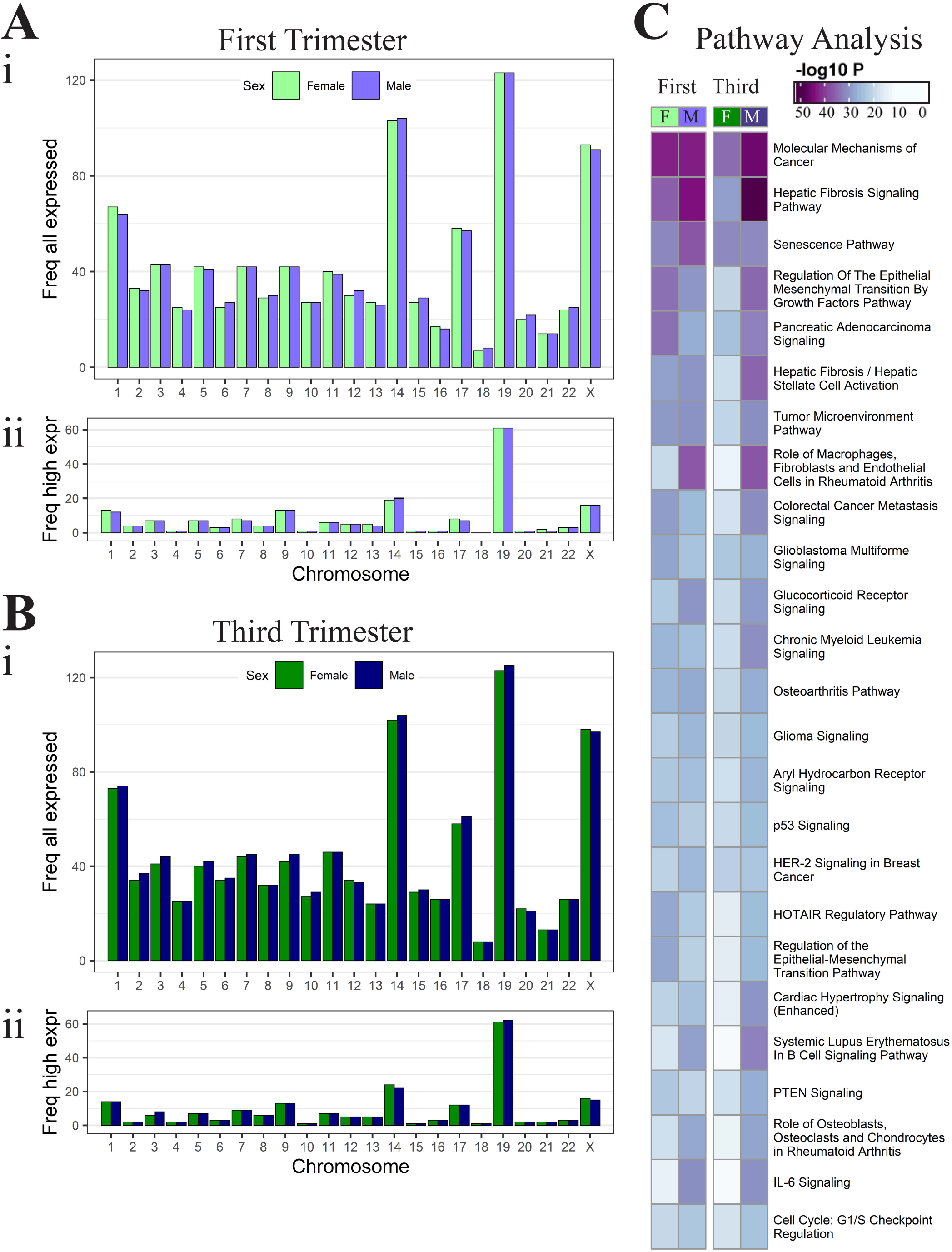
Expressed miRNAs in male and female placentae. (A) Chromosome frequency of miRNAs in male and female placentae at first trimester, (i) all expressed miRNAs (baseMean>10) and (ii) the highest expressed miRNAs (baseMean>2000). Similarly, (B) chromosome frequency at third trimester. (C) Comparison of the pathways targeted by highly expressed miRNAs (baseMean>2000) in first and third trimester female and male placenta.

Third trimester expression was similar with 400 mature miRNAs expressed in the third trimester female placenta derived from 441 precursors (baseMean >10), and 508 mature miRNAs expressed in the third trimester male placenta derived from 570 precursors (baseMean >10, Figure 1Bi) from all 22 autosomes and the X chromosome but not Y. Among the expressed miRNAs, the majority originate with similar sex distribution from chromosome 19 (Female, 123/441 [27.89%]; Male, 125/570 [21.93%]), chromosome 14 (Female, 102/441 [23.13%]; Male, 104/570 [18.25%]), the X chromosome (Female, 98/441 [22.22%]; Male, 97/570 [17.02%]), and chromosome 1 (Female, 73/441 [16.55%]; Male, 74/570 [12.98%]). Among the most highly expressed miRNAs (baseMean >2000), 71 miRNAs were expressed in females and 103 in males, also predominantly from chromosome 19 (Female, 61/71 [85.92%]; Male, 61/103 [59.22%]; Figure 1Bii).

Pathway enrichment analysis was performed using experimentally confirmed targets of the most highly expressed miRNAs. Of the overall most significantly enriched pathways, these were more significant in males compared to females: “Hepatic Fibrosis Signaling” in first and third trimester, “Senescence Pathway” in first trimester placenta, and “Molecular Mechanisms of Cancer” only in third trimester. Of the top 30 significant pathways, the following pathways were more significant in females in the first trimester but became more significant in males in the third trimester: “Regulation of the Epithelial Mesenchymal Transition by Growth Factors”, “Pancreatic Adenocarcinoma Signaling”, “HOTAIR Regulatory Pathway”, and “Regulation of the Epithelial-Mesenchymal Transition” pathway. The following pathways related to immune processes had greater significance in males than females throughout gestation: “Role of Macrophages, Fibroblasts, and Endothelial Cells in Rheumatoid Arthritis”, “Glucocorticoid Receptor Signaling”, “Glioma Signaling”, “Systemic Lupus Erythematosus in B Cell Signaling Pathway”, “Role of Osteoblasts, Osteoclasts, and Chondrocytes in Rheumatoid Arthritis”, “IL-6 Signaling”, and “Neuroinflammation Signaling Pathway” (Figure 1C, Supplemental File 2).

### Differentially expressed miRNAs between male and female placentae

Heatmaps of all expressed miRNAs in the first trimester (Supplemental file 3A) did not demonstrate any clustering of subjects or miRNAs based on sex. Although there were 93 differentially expressed miRNAs, with 57 upregulated in females and 36 upregulated in males (p<0.05, baseMean>10), only 11 miRNAs were differentially expressed following adjustment for multiple comparisons (FDR<0.05, baseMean>10), all upregulated in females (Fig 2Ai). Of the 11 differentially expressed miRNAs (FDR<0.05), the majority had high baseMeans, except miR-429 (female 136.3; male 82.5 baseMeans) and miR-95-3p (female 166.71; male 77.20 baseMeans). The most differentially expressed miRNA was miR-200a-3p with a fold change of 3.49 and baseMeans of 927.5 in females and 525.5 in males. Four of eleven miRNAs were expressed on chromosome X (36.4%), two on chromosome 1 (18.2%), and one each on five other autosomes (9.1% each) (Fig 2Bi).

**Figure 2:**
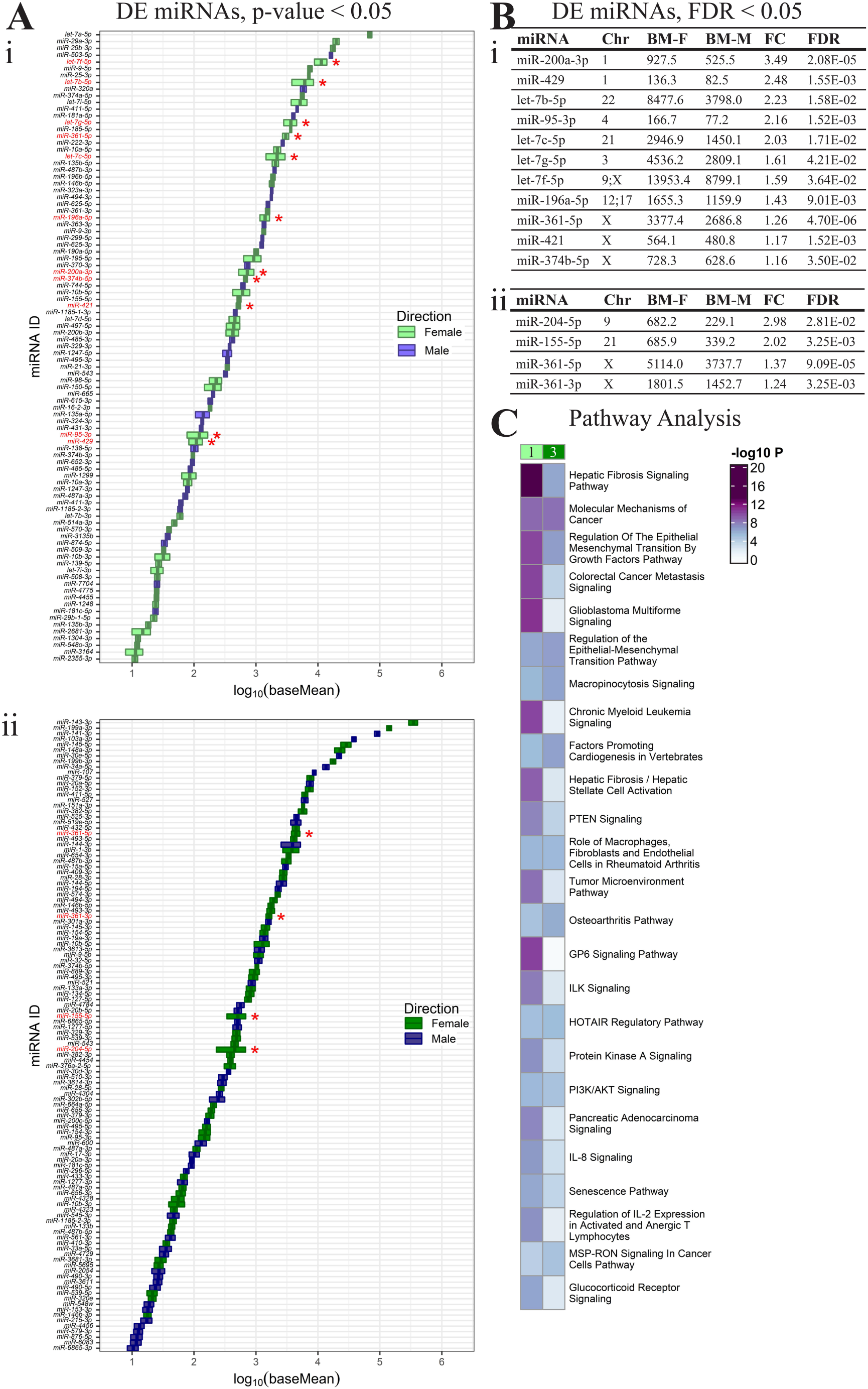
Sexually dimorphic miRNAs in first and third trimester. (A) 93 differentially expressed (DE) miRNAs (57 up in female and 36 in male, baseMean>10, p-value<0.05) listed based on increasing expression (log_10_ baseMean). Red stars and red labels: DE miRNAs that remain significant after multiple comparisons (FDR<0.05) in (i) first and (ii) third trimesters. (B) Significant miRNAs at FDR<0.05 in (i) first and (ii) third trimesters are tabulated with chromosome (Chr) location(s), average baseMeans of female (BM-F) and male (BM-M) samples, fold change (FC, female:male), and FDR. (C) Comparison of the pathways targeted by differentially expressed and female-upregulated miRNAs reveals sexually dimorphic differences between first (1) and third (3) trimester.

In the third trimester, heatmaps of all expressed miRNAs also demonstrated no clustering of subjects or miRNAs based on sex (Supplemental File 3B). Although there were 114 differentially expressed miRNAs, with 64 upregulated in females and 50 upregulated in males (p<0.05, baseMean>10), only 4 miRNAs were differentially expressed following adjustment for multiple comparisons (FDR<0.05, baseMean>10), all upregulated in females (Fig 2Aii). Of the 4 differentially expressed miRNAs (FDR<0.05), the most differentially expressed miRNA was miR-204-5p with 2.98-fold higher expression in females (682.2 female; 229.1 male baseMean). The differentially expressed miRNAs in the third trimester originated from chromosome X (2 miRNAs), 9 (1 miRNA) and 21 (1 miRNA, Fig 2Bii). The miRNA, miR-361-5p was differentially expressed and upregulated in females in both trimesters.

Pathway enrichment analysis was performed using experimentally confirmed targets of the 11 and 4 sexually dimorphic miRNAs in the first and third trimester, respectively. Among the top 10 significant pathways, these were more significantly regulated by sexually dimorphic miRNAs in the first trimester compared to the sexually dimorphic miRNAs in the third trimester: “Hepatic Fibrosis Signaling Pathway”, “Regulation of the Epithelial Mesenchymal Transition by Growth Factors Pathway”, “Colorectal Cancer Metastasis Signaling”, “Glioblastoma Multiforme Signaling”, “Chronic Myeloid Leukemia Signaling”, and “Hepatic Fibrosis / Hepatic Stellate Cell Activation”. The following pathways were regulated by sexually dimorphic miRNAs, with similar significance in first and third trimester: “Molecular Mechanisms of Cancer” and “Role of Macrophages, Fibroblasts, and Endothelial Cells in Rheumatoid Arthritis”. The “GP6 Signaling Pathway” was regulated by sexually dimorphic miRNAs in the first trimester only (Figure 2C, Supplemental File 2).

### Sex-specific differential miRNAs expression across gestation

In order to identify sex specific differences throughout gestation, differential expression analysis of first vs third trimester placenta was performed separately with the female placentae and male placentae. There was significant sample separation by trimester in both sexes (Supplemental File 3C-D). In female placentae, 554 mature miRNAs were significantly differentially expressed between first and third trimester placentae (FDR<0.05), with 301 miRNAs upregulated in the first trimester and 253 miRNAs upregulated in the third trimester placentae (Figure 3Ai-ii). Of the 554 miRNAs derived from 609 precursors in female placenta throughout gestation, the majority originate from chromosomes 19 (first trimester, 34/323 [10.53%]; third trimester, 31/286 [10.84%]), 14, (first trimester, 26/323 [8.05%]; third trimester, 36/286 [12.59%]), and X (first trimester, 33/323 [10.22%]; third trimester, 30/286 [10.49%]) (Figure 3Aii). Of the differentially expressed miRNAs in females across gestation, those with the greatest fold changes (fold change>8) had lower to moderate expression (baseMean 1.18-1261.45) and were primarily elevated in the first trimester, whereas those with the highest expression (baseMean>10,000) had lower fold changes (fold change 0.24-2.15) and were predominantly elevated in the third trimester. The most differentially expressed miRNA was miR-4483, with 42.5-fold higher expression in first trimester placenta (FDR=4.32E-166) and a base mean decrease from 1030.93 to 24.24 in first to third trimester placentae (Figure 3Aiii). In the male cohort, 585 mature miRNAs were significantly differentially expressed between first and third trimester placentae (FDR<0.05), with 309 miRNAs upregulated in the first trimester and 276 miRNAs upregulated in the third trimester placentae (Figure 3Bi-ii).

**Figure 3:**
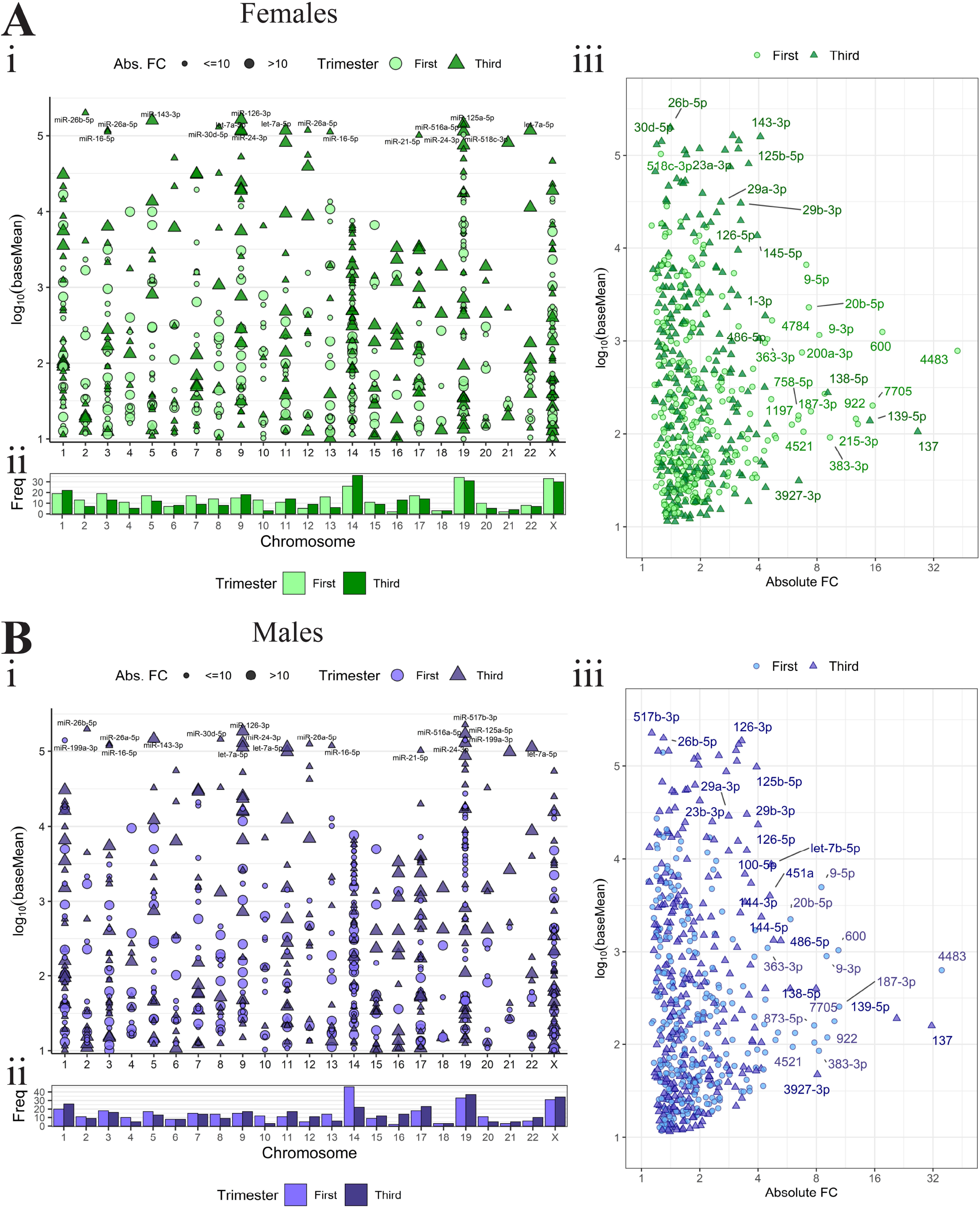
Sex-specific expression differences in miRNAs across gestation. DE miRNAs in females and (B) males. (i) Scatter plot of expression (log_10_ baseMean) distribution across chromosomes for all DE miRNAs at FDR <0.05, baseMean>10. (ii) Chromosome frequency of upregulated miRNAs for each trimester at FDR <0.05, baseMean>10. (iii) Scatter plots of expression (log_10_ baseMean) versus Absolute Fold Change for DE miRNAs (FDR <0.05). For all scatter plots, point shape indicates direction of upregulation.

Chromosomal distribution was similar for males across gestation with 585 miRNAs derived from 651 precursors. The majority of male miRNAs originate from chromosomes 19 (first trimester, 33/332 [9.94%]; third trimester, 39/319 [12.23%]), 14 (first trimester, 46/332 [13.86%]; third trimester, 22/319 [6.89%]), and X (first trimester, 31/332 [9.34%]; third trimester, 34/319 [10.66%]) (Figure 3Bii). Of the differentially expressed miRNAs in males across gestation, those with the greatest fold changes (fold change>8) had lower to moderate expression (baseMean 4.22-835.59) and were primarily elevated in the first trimester, whereas those with the highest expression (baseMean>10,000) had lower fold changes (fold change 0.25-2.25) and were predominantly elevated in the third trimester. The most differentially expressed miRNA was also miR-4483, with 35.5-fold higher expression in first trimester placenta (FDR=1.22E-182) and a baseMean decrease from 936.26 to 26.33 in first to third trimester placentae (Figure 3Biii).

### Common and sex-specific miRNAs differentially expressed across gestation

There were 207 miRNAs differentially expressed (DE) between first and third trimester common to both female and male cohorts. Of these common DE miRNAs, 111 were upregulated in first and 96 were upregulated in third trimester. Common DE miRNAs with strong gestational differences in both sexes (fold change>8 in both) include miR-4483, miR-600, miR-7705, miR-5692b, miR-922, miR-187-3p, miR-383-3p, and miR-9-3p upregulated in first trimester, and miR-137 and miR-139-5p upregulated in third trimester. The common DE miRNA most upregulated in first trimester placenta (compared to third trimester) was miR-4483 with a 42.52-fold difference in female placenta and a 35.48-fold difference in male placenta (Fig. 4Ai). The common DE miRNA most upregulated in third trimester placenta (compared to first trimester) was miR-137 with a 26.49-fold difference in female placenta and a 31.52-fold difference in male placenta (Fig 4Aii).

**Figure 4:**
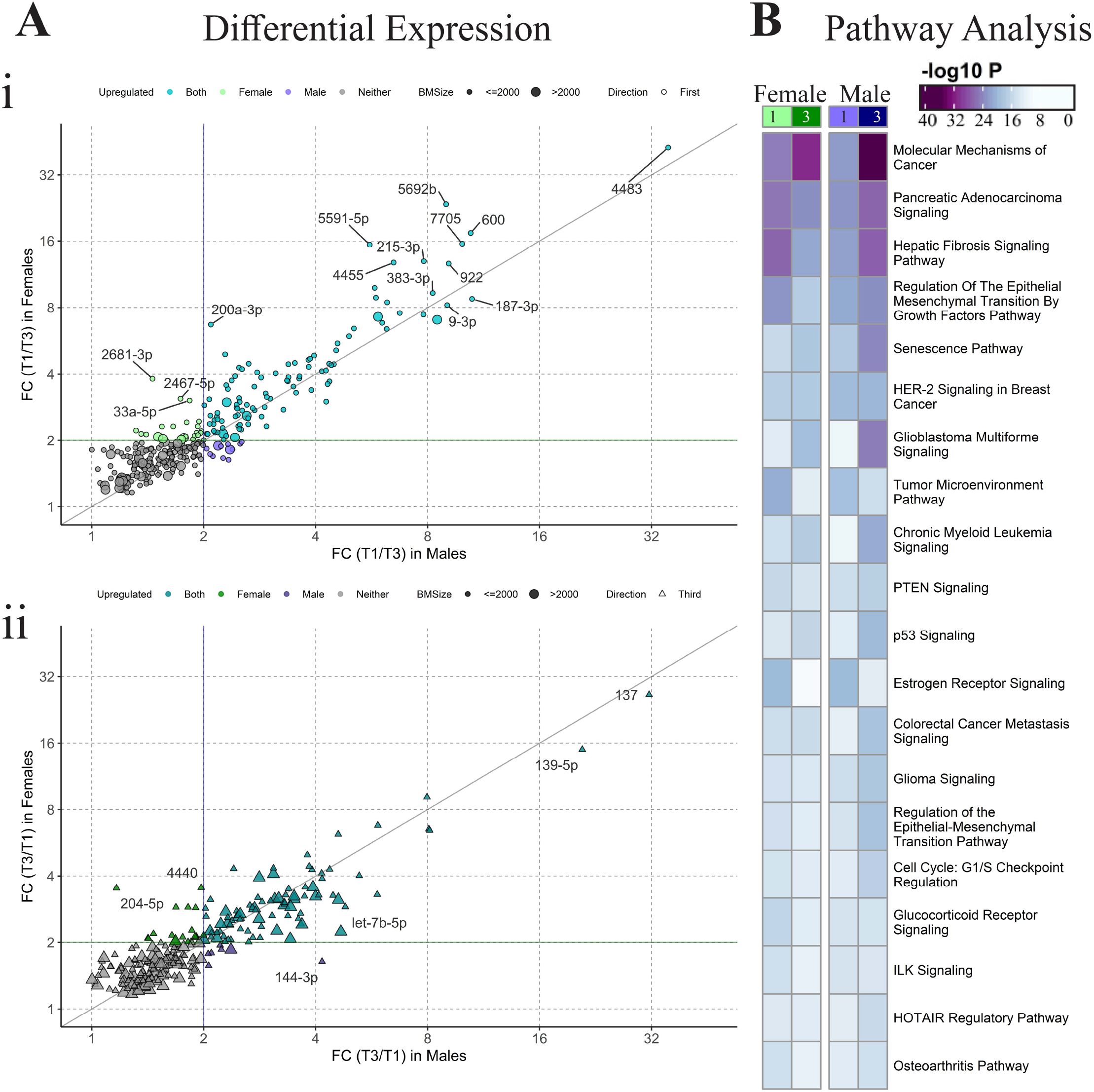
Common and sex-specific miRNA expression changes across gestation. (A) Scatter plots of first vs third trimester DE miRNA fold changes in females compared to males, highlighting miRNAs upregulated in (i) first and (ii) third trimester (FC>1). Colors: common DE miRNAs which reach FC>2 in both sexes (aqua/turquoise), female specific DE miRNAs (green), male specific DE miRNAs (purple), or neither (gray). (B) Comparison of the pathways targeted by the most upregulated miRNAs (FDR<1E-5, baseMean>100 in each group, fold change>2) for each sex cohort in the first vs third trimester analysis.

There was also sex specific differential expression (DE) of miRNAs in each first versus third trimester analysis, suggesting sexually dimorphic miRNA regulation across gestation. The female cohort had more DE miRNAs across gestation that were not present in the male analysis (Fig 4A, Supplemental File 4). There were 44 miRNAs differentially expressed across gestation only in females (FDR<0.05, FC>2, baseMean>10), whereas 21 specifically differentially expressed only in males (FDR<0.05, FC>2, baseMean>10). Of the sex-specific DE miRNAs, female specific DE miRNAs had greater fold changes than male specific DE miRNAs (Fig 4A). Of the female specific DE miRNAs, 23 were upregulated in the first trimester (Fig 4Ai) and 21 were upregulated in the third trimester (Fig. 4Aii). Of female specific DE miRNAs, miR-2681-3p was the most differentially expressed with 3.81-fold higher expression in first trimester placenta (FDR=0.0248) and a baseMean decrease of 20.53 to 2.96 from first to third trimester placentae. Of male specific DE miRNAs, 12 were upregulated in the first trimester (Fig 4Ai) and 9 were upregulated in the third trimester (Fig. 4Aii). Of male specific DE miRNAs, miR-144-3p was the most differentially expressed with 4.16-fold higher expression in third trimester placenta (FDR=3.75E-16) and a baseMean increase of 1142.11 to 4752.80 from first to third trimester placentae.

Pathway enrichment analysis on experimentally confirmed targets of the sex-specific DE miRNAs across gestation identified “Molecular Mechanisms of Cancer” to be most significant in the third trimester for both sexes, with greater significance in males. The “Senescence Pathway” was also more significant in the third trimester for both sexes. Top pathways more significant in first versus third in female placentae include “Pancreatic Adenocarcinoma Signaling”, “Hepatic Fibrosis Signaling”, and “Regulation of the Epithelial Mesenchymal Transition by Growth Factors”, whereas these pathways were more significant in third versus first trimester in males (Figure 4B, Supplemental File 2).

### Sex differences in the placenta-specific C14MC and C19MC Clusters

Among the DE miRNAs across gestation, in the C14MC cluster, miR-1197 was the most differentially expressed in both sexes and upregulated in the first trimester. In the C19MC cluster, miR-520c-3p was the most differentially expressed in both sexes and upregulated in the third trimester (Figure 5Ai-ii). Of the C14MC and C19MC, there were 2 that were differentially expressed across gestation in females but not males (miR-654-5p up in first (C14MC) and miR-541-3p up in third trimester (C14MC)). Three miRNAs were differentially expressed across gestation in males but not females (miR-376a-3p, miR-376a-5p, and miR-476b-3p, all upregulated in third trimester (C14MC)) (Supplementary File 4). Among the DE miRNAs across gestation, the female cohort had 104 DE miRNAs on C14MC (37 upregulated in the first trimester and 55 in the third trimester) and 113 DE miRNAs on C19MC (73 upregulated in the first trimester and 36 in the third trimester, FDR<0.05, baseMean>10, Ref barplot). In the male cohort, there were 102 DE miRNAs on C14MC (62 upregulated in the first trimester and 30 upregulated in third trimester) and 113 DE miRNAs on C19MC (55 upregulated in the first trimester and 54 in third trimester, FDR<0.05, baseMean>10, Figure 5Bi-ii). In females, more DE miRNAs were upregulated in the third trimester from C14MC compared to the first trimester, whereas there were more differentially expressed miRNAs that were upregulated from C19MC in the first trimester, suggesting these clusters play different roles throughout gestation in females, with C14MC having a greater regulatory role in the third trimester and C19MC a greater regulatory role in the first trimester. In contrast, there were more differentially expressed miRNAs that were upregulated from C14MC in the first trimester male placenta compared to the third trimester, suggesting a greater regulatory role of C14MC in males in the first trimester. (Figure 5Bi-ii).

**Figure 5:**
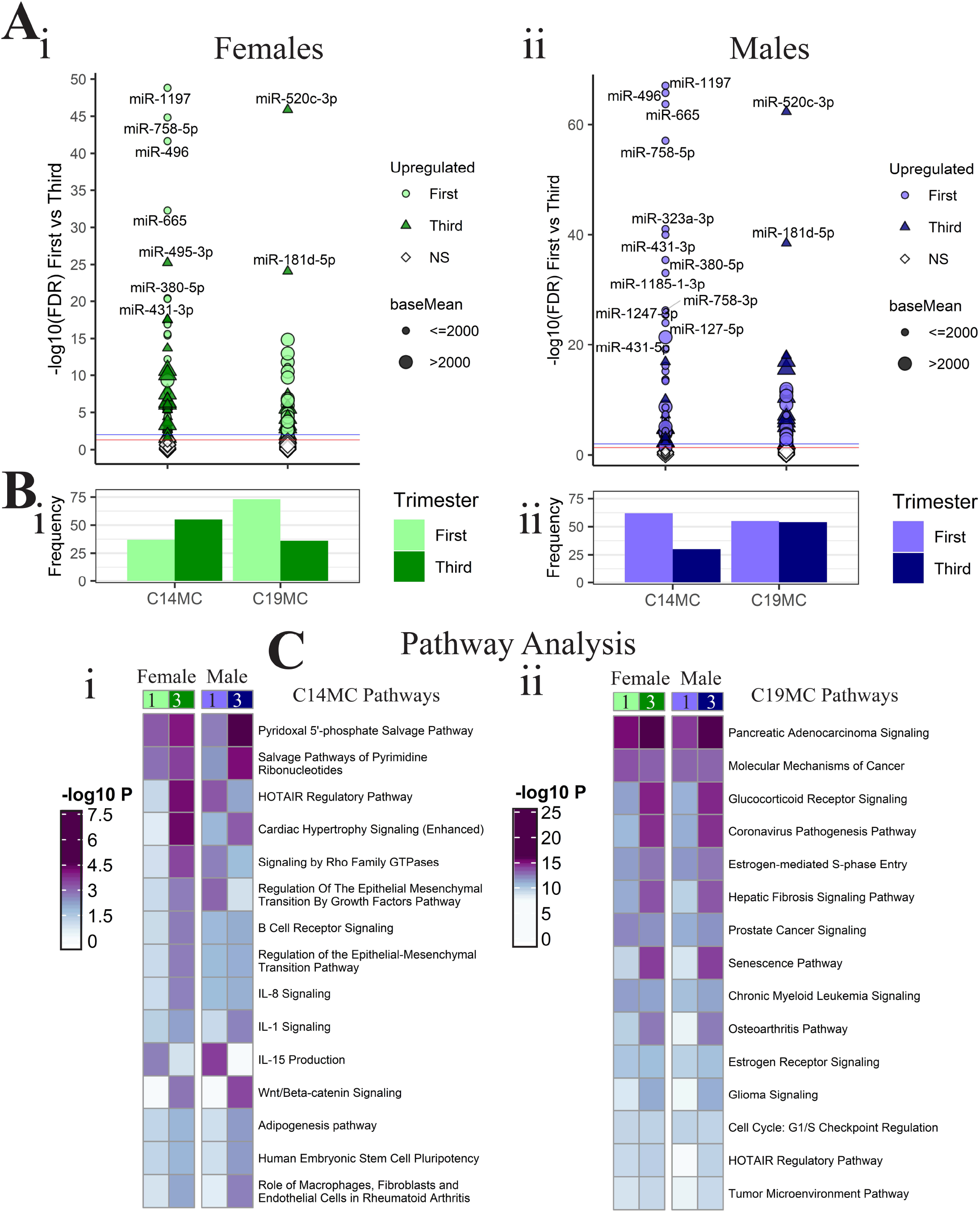
Expression differences in C14MC and C19MC clusters. Scatter plots for (Ai) female and (Bi) male cohorts show miRNAs that are upregulated in first or third trimester (color/shape) and highly expressed (size) for each cluster. (Aii-Bii) Corresponding frequency plots count the upregulated miRNAs for each trimester (color). (Ci-ii) Comparison of the pathways targeted by these upregulated miRNAs, by cluster.

Pathway enrichment analysis on experimentally confirmed targets of the C14MC miRNAs identified “Pyridoxal 5’-Phosphate Salvage Pathway”, “Salvage Pathways of Pyrimidine Ribonucleotides”, “Cardiac Hypertrophy Signaling (Enhanced)”, “IL-1 Signaling”, and “Wnt/β-Catenin-Signaling” to be the most highly significant in the third trimester for both sexes. “HOTAIR Regulatory Pathway”, “Signaling by Rho Family GTPases”, and “Regulation of the Epithelial Mesenchymal Transition by Growth Factors” were highly significant in females in the third trimester but highly significant in males in the first trimester, with the latter not significant in first trimester females and third trimester males. “IL-15 Production” was different from all previous patterns in that it was highly significant in first trimester for both sexes. The C19MC pathways were less sexually dimorphic across gestation with most of the pathways more significant in third versus first trimester. Of the top 30, no pathways were highly significant in first trimester compared to third, although a few were similar (Figure 5C, Supplemental File 2).

## Discussion

We identified the sex specific microRNA signature of the first and third trimester placenta in pregnancies resulting in live births. There were similar numbers of miRNAs expressed in both sexes in each trimester, with similar chromosome distributions and peaks at chromosomes 1, 14, 19, and X as previously described [19, 54-56]. Overall, the majority of miRNA were not sexually dimorphic with 11 differentially expressed miRNAs following multiple comparisons in first and 4 in third trimester placentae, all upregulated in females. Sex specific expression differences across gestation were similar, with similar chromosome peaks, however chromosomal distributions were sexually dimorphic in chromosome 14 and 19, with chromosome 14 more representative in the third trimester in females but more representative in the first trimester in males, likely due to C14MC and C19MC. There were first versus third trimester DE miRNAs common to females and males, including miR-4483 which was upregulated in the first trimester and miR-137 and miR-139-5p up in third trimester. There were twice as many female-specific gestational differences as male-specific (44 miRNAs vs 21 miRNAs), indicating that miRNA abundance across human gestation is sexually dimorphic. To our knowledge, this is the largest normative sex dimorphic and sex specific placenta miRNA atlas across gestation.

Although there were similar numbers of miRNAs expressed in both sexes in each trimester, with similar chromosome distributions, immune mediated pathways had greater significance in males in first and third trimester placenta compared to females. miRNAs have been identified in regulating immune function, which may be sexually dimorphic [57] that may also translate at the placenta, and these sexually dimorphic immune mediated pathways may impact placental function including malperfusion and subsequent sequela [58]. Furthermore, this altered intrauterine environment may have long term impact, including deficient development of the immune system into childhood [59-61], predispose to inflammatory and immune disease states that develop in adulthood [61-63], including metabolic syndrome and cardiovascular disease in adulthood [64-69], which are sexually dimorphic disease states.

Sex differences in the first and third trimester were small, but differences were more prevalent in the first trimester placenta, 11 sexually dimorphic miRNAs in the first trimester and 4 in the third trimester, all upregulated in female placentae compared to male placentae. Greater sexual dimorphism in miRNAs may also account for more significant upregulated pathways identified in the first compared to the third trimester placenta, suggesting miRNAs have an impact on placental development early in gestation in a sex specific manner, a result consistent with previous studies [10, 21, 70]. The only miRNA that was sexually dimorphic in the first and third trimester was miR-361-5p. This miRNA has been minimally studied, with no references to placental development, however, Tsamou *et al*. report that this miRNA was also sexually dimorphic in newborns, with higher expression in girls [71].

Although other studies have identified sexually dimorphic miRNA expression in the placenta, these studies were small, which may lead to subject variability [72], or utilized microarrays [71] and most of them were in placenta disease states where differences may be attributed to development of placental dysfunction which may be the result of sexually dimorphic miRNA regulation of these sexually dimorphic disease states early in gestation [10, 73]. Recently, sexually dimorphic miRNA expression was described by Eaves in second trimester placenta of extremely low gestational age newborns. Of those identified by Eaves, miR-361-5p, miR-196a-5p, and miR-374b-5p were also sexually dimorphic in the first trimester in our study [20]. These miRNA may ultimately be sex specific early markers of extremely low gestational age. Another recent study by Guo *et al*. of a small number of first and third trimester sexually dimorphic samples also found miR-361-3p and miR-361-5p in common with our third trimester samples [72], suggesting these may be specific markers of normal gestational age. In addition, other miRNAs defined in our study have been previously shown to be associated with preterm labor, such as miR-421, miR-374b-5p, and miR-155-5p [74]. Other gestational disease states have been previously studied as well, including recurrent spontaneous abortion, associated with miR-196a-5p [75], and pre-eclampsia, associated with miR-374b-5p [30], and may ultimately become sex specific markers of these sex dimorphic placental pathologic states.

There were 207 miRNAs with differential expression across gestation common in both sexes. These are potentially critical miRNAs, particularly those that are expressed early in gestation, that can be used for future biomarker development, since they are independent of fetal sex. miR-4483, which is upregulated in first compared to third trimester, was also found to be strongly downregulated in the second trimester and hence likely plays a significant role in the first trimester placenta that is sex independent [10]. There were also sex specific miRNAs that had differential expression across gestation, suggesting miRNA regulation of placental development throughout gestation is sexually dimorphic. These miRNAs are important for sex specific normal placental development and may provide a foundation for miRNA expression changes leading to placental pathology in a sex specific fashion.

In the placenta-specific C14MC and C19MC[22, 23] none of the miRNAs were sexually dimorphic in the first or third trimester. However, there were sex specific differences across gestation in C14MC and C19MC. In females, there were more differentially expressed miRNAs that were upregulated in the third trimester from C14MC compared to the first trimester, whereas there were more differentially expressed miRNAs that were upregulated from C19MC in the first trimester, suggesting these clusters play different roles throughout gestation in females, with C14MC having a greater regulatory role in the third and C19MC in the first trimester. In contrast, in males, there were more differentially expressed miRNAs that were upregulated from C14MC in the first compared to the third trimester, suggesting a greater regulatory role of C14MC in males in the first trimester. There were two female specific differentially expressed miRNAs across gestation, miR-541-3p upregulated in third trimester, and miR-654-5p upregulated in first trimester, both from C14MC. There were three male specific differentially expressed miRNAs across gestation from the miR-376 precursor, miR-376a-3p, miR-376a-5p, and miR-376b-3p, all upregulated in third trimester. All of these miRNAs have been associated with cancer including reproductive cancers [76-79], which may be suggestive that these miRNAs are under sex hormone control, which is fetal sex specific early in gestation during fetal sex differentiation [80, 81]. The sex specific differences across gestation in C14MC and C19MC may account for differences in the identified canonical pathways. “HOTAIR Regulatory Pathway”, “Signaling by Rho Family GTPases”, and “Regulation of the Epithelial Mesenchymal Transition by Growth Factors” were highly significant in females in the third trimester but highly significant in males in the first trimester, with the latter not significant in first trimester females and third trimester males.

The major strength of this study is the cohort size of normal healthy pregnancies resulting in delivery and the availability of detailed demographic information including birth outcomes. Additionally, the use of high-throughput sequencing, as opposed to other techniques such as arrays, allows for greater confidence regarding differential expression, since all known miRNA species previously annotated in the human genome are considered. Although other studies have identified sexually dimorphic miRNA expression in the placenta, these studies were small, [72] and most of them were in placenta disease states where differences may be attributed to development of placental dysfunction which may be the result of sexually dimorphic miRNA regulation of these sexually dimorphic disease states [71, 73].

Our study has some limitations. Female and male singleton pregnancies in the first and third trimester were matched for maternal age, race, ethnicity and pre-existing medical conditions, however, when comparing sexes across gestation, differences in parental and fetal race and ethnicity although small were identified. Maternal pre-existing medical conditions were not significantly different among the groups but there were a few mothers that developed hypertension in the third trimester cohorts. However, we did not control for outcomes since subjects were enrolled in the first trimester. Pre-pregnancy BMI was greater in mothers of males in the third trimester cohort compared to the first trimester cohort. Although sexual dimorphism in miR-210 expression has been identified in the placenta associated with maternal obesity [82], our cohorts were not obese and BMI differences were only identified in the male specific cohorts. As a result, miR-210 was not identified to be different among the groups.

Overall, our goals were: first, to identify the normative sex dimorphic miRNA signatures in first and third trimester and second, identify and compare the normative sex stable and sex specific normative miRNA signature across gestation in a healthy population. With the identification of these normative signatures, future studies on placental pathologic disease states can be performed, controlling for variability due to fetal sex and gestational age. Furthermore, due to their small size and stability, these miRNAs may become potential biomarkers of disease early in gestation, particularly miRNAs from plasma exosomes,[83-85] and utilized as additional metadata for machine learning models of placental disease [86].

## Acknowledgements

The authors would like to acknowledge the members within the department of OBGYN who assisted with subject recruitment, and most of all we are grateful for the subjects who are willing to participate in our studies.

## Conflict of Interest

None to declare.

## Author Contributions

Author(s) contributed to conceptualization and design of work (AEF, TLG, NVJ, LEE, KL, CJ, YZ, YA, HRT, JW III, MDP), acquisition of samples (ELC, RAB, ES, RD, JW III), sample processing (TLG, NVJ, ELC, RD, YL, CS, BL, TS), analysis (AEF, TLG, NVJ, DW, YW, JT, JLC, ETW, MDP), and interpretation of data (AEF, TLG, NVJ, DW, YW, MDP). The original draft was written by AEF, TLG, NVJ, LEE, and MDP.

## Supplemental Information

**Supplemental File 1: Principal Components Analysis**. 1.1ABC: Female vs Male principal components graphs of (A)PC1 vs PC2, (B)PC1 vs PC3, and (C)PC2 vs PC3 for first trimester samples. 1.2ABC: Female vs Male principal components graphs of (A)PC1 vs PC2, (B)PC1 vs PC3, and (C)PC2 vs PC3 for third trimester samples. 1.3ABC: First v Third principal components graphs of (A)PC1 vs PC2, (B)PC1 vs PC3, and (C)PC2 vs PC3 for female samples. 1.4ABC: First v Third principal components graphs of (A)PC1 vs PC2, (B)PC1 vs PC3, and (C)PC2 vs PC3 for male samples. [.pdf file]

**Supplemental File 2:** Compilation of full target gene enrichment analysis results from IPA Core Analysis. [Excel .xlsx file]

**Supplemental File 3: Heatmaps of Subjects vs miRNAs**. Heatmaps depicting sex-differential data in (Ai) first and (Aii) third trimester and sex-specific data in (Bi) females and (Bii) males. All heatmaps restricted to miRNAs of baseMean>10. [.pdf file]

**Supplemental File 4**: Compilation of miRNA expression analysis results. [Excel .xlsx file]

## Notes

**Grant Support:** This work was supported by the National Institute of Health grants: R01 HD091773, R01 HD074368, T32 DK007770, U01 EB026421, and R01 AI154535. The funding agency was not involved in the design, analysis, or interpretation of the data reported. The content is solely the responsibility of the authors and does not necessarily represent the official views of the National Institutes of Health.

### Competing Interest Statement

The authors have declared no competing interest.

## References

1. Broere-Brown ZA, Adank MC, Benschop L, Tielemans M, Muka T, Gonçalves R, Bramer WM, Schoufour JD, Voortman T, Steegers EAP, Franco OH, Schalekamp-Timmermans S. Fetal Sex and Maternal Pregnancy Outcomes: A Systematic Review and Meta-Analysis. Biology of Sex Differences 2020; 11.

2. Lorente-Pozo S, Parra-Llorca A, Torres B, Torres-Cuevas I, Nuñez-Ramiro A, Cernada M, García-Robles A, Vento M. Influence of Sex on Gestational Complications, Fetal-to-Neonatal Transition, and Postnatal Adaptation. Frontiers in Pediatrics 2018; 6.

3. Traccis F, Frau R, Melis M. Gender Differences in the Outcome of Offspring Prenatally Exposed to Drugs of Abuse. Frontiers in Behavioral Neuroscience 2020; 12.

4. Sheiner E. The Relationship Between Fetal Gender and Pregnancy Outcomes. Archives of Gynecology and Obstetrics 2007; 275:317–319.

5. Kiserud T, Benachi A, Hecher K, Perez RG, Carvalho J, Piaggio G, Platt LD. The World Health Organization Fetal Growth Charts: Concept, Findings, Interpretation, and Application. American Journal of Obstetrics and Gynecology 2018; 218:S619–S629.

6. Basso O, Olsen J. Sex Ratio and Twinning in Women with Hyperemesis or Pre-Eclamsia. Epidemiology 2001; 12:747–749.

7. Kuru O, Sen S, Akbayir O, Goksedef BPC, Ozsurmeli M, Attar E, Saygili H. Outcomes of Pregnancies Complicated by Hyperemesis Gravidarum. Archives of Gynecology and Obstetrics 2011; 285:1517–1521.

8. Nurmi M, Rautava P, Gissler M, Vahlberg T, Polo-Kantola P. Recurrence Patterns of Hyperemesis Gravidarum. American Journal of Obstetrics and Gynecology 2018; 219:e1–10.

9. Boss AL, Chamley LW, James JL. Placental Formation in Early Pregnancy: How is the Center of the Placenta Made. Human Reproduction Update 2018; 24:750–760.

10. Gonzalez TL, Sun T, Koeppel AF, Lee B, Wang ET, Farber CR, Rich SS, Sundheimer LW, Buttle RA, Chen Y-DI, Rotter JI, Turner SD, et al. Sex Differences in the Late First Trimester Human Placenta Transcriptome. Biology of Sex Differences 2018; 9.

11. Buckberry S, Bianco-Miotto T, Bent SJ, Dekker GA, Roberts CT. Integrative Transcriptome Meta-Analysis Reveals Widespread Sex-Biased Gene Expression at the Human Fetal-Maternal Interface. Molecular Human Reproduction 2014; 20.

12. Hayder H, O’Brien J, Nadeem U, Peng C. MicroRNAs: Crucial Regulators of Placental Development. Reproduction 2018; 155:R259–R271.

13. Mouillet J-F, Ouyang Y, Coyne C, Sadovsky Y. MicroRNAs in placental health and disease. American Journal of Obstetrics and Gynecology 2015; 213:S163–S172.

14. de Sousa MC, Gjorgjieva M, Dolicka D, Sobolewski C, Foti M. Deciphering miRNAs’ Action through miRNA editing. International Journal of Molecular Sciences 2019; 20.

15. Ali A, Bouma GJ, Anthony RV, Winger QA. The Role of LIN28-let-7-ARID3B Pathway in Placental Development. International Journal of Molecular Sciences 2020; 21.

16. Zhang WM, L Xin PC, Zhang Y, Liu Z, Yao N, Ma Y-Y. Effect of miR-133 on Apoptosis of Trophoblasts in Human Placenta Tissues via Rho/ROCK signaling pathway. European Review for Medical and Pharmacological Sciences 2019; 23:10600–10608.

17. Zhang WM, Liu X-L, Yang Y, Ayi N, Li L-H. Highly Expressed microRNA-940 Promotes Early Abortion by Regulating Placenta Implantation by Targeting ZNF672. European Review for Medical and Pharmacological Sciences 2019; 23:2693–2700.

18. Burnett LA, Nowak RA. Exosomes mediate embryo and maternal interactions at implantation and during pregnancy. Frontiers in Bioscience (Scholar Edition) 2016; 8:79–96.

19. Gonzalez TL, Eisman LE, Joshi NV, Flowers AE, Wu D, Wang Y, Santiskulvong C, Tang J, Buttle RA, Sauro E, Clark EL, DiPentino R, et al. High-throughput miRNA-sequencing of the human placenta: expression throughout gestation. bioRxiv 2021:2021.2002.2004.429392.

20. Eaves LA, Phookphan P, Rager JE, Bangma J, Santos Jr HP, Smeester L, O’Shea TM, Fry RC. A role for microRNAs in the epigenetic control of sexually dimorphic gene expression in the human placenta. Epigenomics 2020; Epub ahead of Print:1-16.

21. Gu Y, Sun J, Groome L, Wang Y. Differential miRNA Expression Profiles Between the First and Third Trimester Human Placentas. American Journal of Physiology: Endocrinology and Metabolism 2013; 304:E836–E843.

22. Bentwich I, Avniel A, Karov Y, Aharonov R, Gilad S, Barad O, Barzilai A, Einat P, Einav U, Meiri E, Sharon E, Spector Y, et al. Identification of hundreds of conserved and nonconserved human microRNAs. Nat Genet 2005; 37:766–770.

23. Morales-Prieto DM, Ospina-Prieto S, Chaiwangyen W, Schoenleben M, Markert UR. Pregnancy-associated miRNA-clusters. J Reprod Immunol 2013; 97:51–61.

24. Liang Y, Ridzon D, Wong L, Chen C. Characterization of microRNA expression profiles in normal human tissues. BMC Genomics 2007; 8:166.

25. Jinesh GG, Flores ER, Brohl AS. Chromosome 19 miRNA cluster and CEBPB expression specifically mark and potentially drive triple negative breast cancers. PLoS One 2018; 13:e0206008.

26. Nguyen PN, Huang CJ, Sugii S, Cheong SK, Choo KB. Selective activation of miRNAs of the primate-specific chromosome 19 miRNA cluster (C19MC) in cancer and stem cells and possible contribution to regulation of apoptosis. J Biomed Sci 2017; 24:20.

27. Radovich M, Solzak JP, Hancock BA, Conces ML, Atale R, Porter RF, Zhu J, Glasscock J, Kesler KA, Badve SS, Schneider BP, Loehrer PJ. A large microRNA cluster on chromosome 19 is a transcriptional hallmark of WHO type A and AB thymomas. Br J Cancer 2016; 114:477–484.

28. Noguer-Dance M, Abu-Amero S, Al-Khtib M, Lefevre A, Coullin P, Moore GE, Cavaille J. The primate-specific microRNA gene cluster (C19MC) is imprinted in the placenta. Hum Mol Genet 2010; 19:3566–3582.

29. Dunham A, Matthews LH, Burton J, Ashurst JL, Howe KL, Ashcroft KJ, Beare DM, Burford DC, Hunt SE, Griffiths-Jones S, Jones MC, Keenan SJ, et al. The DNA sequence and analysis of human chromosome 13. Nature 2004; 428:522–528.

30. Zhu X-M, Han T, Sargent IL, Yin G-W, Yao Y-Q. Differential expression profile of microRNAs in human placentas from preeclamptic pregnancies vs normal pregnancies. American Journal of Obstetrics and Gynecology 2009; 200:661.e661-661.e667.

31. Ishibashi O, Ohkuchi A, Ali M, Kurashina R, Luo S-S, Ishikawa T, Takizawa T, Hirashima C, Takahashi K, Migita M, Ishikawa G, Yoneyama K, et al. Hydroxysteroid (17-β) Dehydrogenase 1 is dysregulated by Mir-210 and Mir-518c that are aberrantly expressed in preeclamptic placentas: A novel marker for predicting preeclampsia. Hypertension 2012; 59:265–273.

32. Yang S, Li H, Ge Q, Guo L, Chen P. Deregulated microRNA species in the plasma and placenta of patients with preeclampsia. Molecular Medicine Reports 2015; 12:527–534.

33. Anton L, Olaerin-George AO, Hogenesch JB, Elovitz MA. Placental expression of miR-517a/b and miR-517c contributes to trophoblast dysfunction and preeclampsia. PLoS One 2015; 10.

34. Vashukova ES, Glotov AS, Fedotov PV, Efimova OA, Pakin VS, Mozgavaya EV, Pendina AA, Tikhonov AV, Koltsova AS, Baranov VS. Placental microRNA expression in pregnancies complicated by superimposed pre-eclampsia on chronic hypertension. Molecular Medicine Reports 2016; 14:22–32.

35. Hromadnikova I, Kotlabova K, Ondrackova M, Pirkova P, Kestlerova A, Novotna V, Hympanova L, Krofta L. Expression profile of C19MC microRNAs in placental tissue in pregnancy-related complications. DNA and Cell Biology 2015; 34:437–457.

36. Sõber S, Rull K, Reiman M, Ilisson P, Mattila P, Laan M. RNA sequencing of chorionic villi from recurrent pregnancy loss patients reveals impaired function of basic nuclear and cellular machinery. Scientific Reports 2016; 6.

37. Higashijima A, Miura K, Mishima H, Kinoshita A, Jo O, Abe S, Hasegawa S, Miura S, Yamasaki K, Yashida A, Yashiura K-I, Masuzaki H. Characterization of placenta-specific microRNAs in fetal growth restriction pregnancy. Prenatal Diagnosis 2013; 33:214–222.

38. Herbert JF, Millar JA, Raghavan R, Romney A, Podrabsky JE, Rennie MY, Felker AM, O’Tierney-Ginn P, Morita M, DuPriest EA, Morgan TK. Male fetal sex affects uteroplacental angiogenesis in growth restriction mouse model. Biology of Reproduction 2021; 104:924–934.

39. Li J, Song L, Zhou L, Wu J, Sheng C, Chen H, Liu Y, Gao S, Huang W. A microRNA signature in gestational diabetes mellitus associated with risk of macrosomia. Cellular Physiology and Biochemistry 2016; 37:243–252.

40. Ding R, Guo F, Zhang Y, Liu X-M, Xiang Y-Q, Zhang C, Liu Z-W, Sheng J-Z, Huang H-F, Zhang J-Y, Fan J-X. Integrated Transcriptome Sequencing Analysis Reveals Role of miR-138-5p/TBL1X in Placenta from Gestational Diabetes Mellitus. Cellular Physiology and Biochemistry 2018; 51:630–946.

41. Pisarska MD, Akhlaghpaur M, Lee B, Barlow GM, Xu N, Wang ET, Mackey AJ, Farber CR, Rich SS, Rotter JI, Chen YI, Goodorzi MO, et al. Optimization of techniques for multiple platform testing in small precious samples such as human chorionic villus sampling. Prenatal Diagnosis 2016; 36:1061–1070.

42. Martin M. Cutadapt removes adapter sequences from high-throughput sequencing reads. EMBnet Journal 2011; 17.

43. Langmead B, Trapnell C, Pop M, Salzberg SL. Ultrafast and memory-efficient alignment of short DNA sequences to the human genome. Genome Biology 2009; 10.

44. Kozomara A, Griffiths-Jones S. miRBase: annotating high confidence microRNAs using deep sequencing data. Nucleic Acids Research 2014; 42:D68–D73.

45. Love MI, Huber W, Anders S. Moderated estimation of fold change and dispersion for RNA-seq data with DESeq2. Genome Biology 2014; 15.

46. Durinck S, Moreau Y, Kasprzyk A, Davis S, De Moor B, Brazma A, Huber W. BioMart and Bioconductor: a powerful link between biological databases and microarray data analysis. Bioinformatics 2005; 21:3439–3440.

47. Durinck S, Spellman PT, Birney E, Huber W. Mapping identifiers for the integration of genomic datasets with the R/Bioconductor package biomaRt. Nature Protocols 2009; 4:1184–1191.

48. Krämer A, Green J, Pollard JJ, Tugendreich S. Causal analysis approaches in Ingenuity Pathway Analysis. Bioinformatics 2014; 30:523–530.

49. Xiao F, Zuo Z, Cai G, Kang S, Gao X, Li T. miRecords: an integrated resource for microRNA-target interactions. Nucleic Acids Research 2009; 37:D105–110.

50. Karagkouni D, Paraskevopoulou MD, Chatzopoulos S, Vlachos IS, Tastsoglou S, Kanellos I, Papadimitriou D, Kavakiotis I, Maniou S, Skoufous G, Vergoulis T, Dalamagas T, et al. DIANA-TarBase v8: a decade-long collection of experimentally supported miRNA-gene interactions. Nucleic Acids Research 2018; 46:D239–D246.

51. Lewis BP, Jones-Rhoades MW, Bartel DP, Burge CP. Prediction of mammalian microRNA targets. Cell 2003; 115:787–798.

52. Lee B, Kroener LL, Xu N, Wang ET, Banks A, Williams III J, Goodarzi MO, Chen Y-DI, Tang J, Wang Y, Gangalapudi V, Pisarska MD. Function and Hormonal Regulation of GATA3 in Human First Trimester Placentation. Biology of Reproduction 2016; 95:1–9.

53. Xu N, Barlow GM, Cui J, Wang ET, Lee B, Akhlaghpaur M, Kroener L, Williams III J, Rotter JI, Chen Y-DI, Goodarzi MO, Pisarska MD. Comparison of Genome-Wide and Gene-Specific DNA Methylation Profiling in First-Trimester Chorionic Villi From Pregnancies Conceived with Infertility Treatments. Reproductive Sciences 2017; 24:996–1004.

54. Liang Y, Ridzon D, Wong L, Chen C. Characterization of microRNA expression profiles in normal human tissues. BMC Genomics 2007; 8.

55. Noguer-Dance M, Abu-Amero S, Al-Khtib M, Lefevre A, Coullin P, Moore GE, Cavaille J. The primate-specific microRNA gene cluster (C19MC) is imprinted in the placenta. Human Molecular Genetics 2010; 19:3566–3582.

56. Morales-Prieto DM, Ospina-Prieto S, Chaiwangyen W, Schoenleben M, Markert UR. Pregnancy-associated miRNA-clusters. Journal of Reproductive Immunology 2013; 97:51–61.

57. Dai R, Ahmed SA. Sexual dimorphism of miRNA expression: a new perspective in understanding sex bias of autoimmune diseases. Therapeutics and Clinical Risk Management 2014; 10:151–163.

58. Eloundou SN, Lee J, Wu D, Lei J, Feller MC, Ozen M, Zhu Y, Hwang M, Jia B, Zie H, Clemens JL, McLane MW, et al. Placental malperfusion in response to intrauterine inflammation and its connection to fetal sequelae. PLoS One 2019; 14.

59. Ege MJ, Bieli C, Frei R, van Strien RT, Riedler J, Ublagger E, Schram-Bijkerk D, Brunekreef B, van Hage M, Scheynius A, Pershagen G, Benz MR, et al. Prenatal farm exposure is related to the expression of receptors of the innate immunity and to atopic sensitization in school-age children. J Allergy Clin Immunol 2006; 117:817–823.

60. Schaub B, Liu J, Hoppler S, Schleich I, Huehn J, Olek S, Wieczorek G, Illi S, von Mutius E. Maternal farm exposure modulates neonatal immune mechanisms through regulatory T cells. J Allergy Clin Immunol 2009; 123:774-782.e775.

61. Chen T, Liu HX, Yan HY, Wu DM, Ping J. Developmental origins of inflammatory and immune diseases. Mol Hum Reprod 2016; 22:858–865.

62. Huizink AC, Mulder EJ, Buitelaar JK. Prenatal stress and risk for psychopathology: specific effects or induction of general susceptibility? Psychol Bull 2004; 130:115–142.

63. Fowden AL, Giussani DA, Forhead AJ. Intrauterine programming of physiological systems: causes and consequences. Physiology (Bethesda) 2006; 21:29–37.

64. Barker DJ, Eriksson JG, Forsen T, Osmond C. Fetal origins of adult disease: strength of effects and biological basis. Int J Epidemiol 2002; 31:1235–1239.

65. Barker DJ, Osmond C, Law CM. The intrauterine and early postnatal origins of cardiovascular disease and chronic bronchitis. J Epidemiol Community Health 1989; 43:237–240.

66. Barker DJ, Winter PD, Osmond C, Margetts B, Simmonds SJ. Weight in infancy and death from ischaemic heart disease. Lancet 1989; 2:577–580.

67. Barker DJ, Hales CN, Fall CH, Osmond C, Phipps K, Clark PM. Type 2 (non-insulin-dependent) diabetes mellitus hypertension and hyperlipidaemia (syndrome X) relation to reduced fetal growth. Diabetologia 1993; 36:62–67.

68. Barker DJ, Hales CN, Fall CH, Osmond C, Phipps K, Clark PM. Type 2 (non-insulin-dependent) diabetes mellitus, hypertension and hyperlipidaemia (syndrome X): relation to reduced fetal growth. Diabetologia 1993; 36:62–67.

69. Vaughan OR, Rosario FJ, Powell TL, Jansson T. Regulation of Placental Amino Acid Transport and Fetal Growth. Prog Mol Biol Transl Sci 2017; 145:217–251.

70. Sun T, Gonzalez TL, Deng N, DiPentino R, Clark EL, Lee B, Tang J, Wang Y, Stripp BR, Yao C, Tseng H-R, Karumanchi SA, et al. Sexually dimorphic crosstalk at the maternal-fetal interface. Journal of Clinical Endocrinology and Metabolism 2020; 105:e4831–e4847.

71. Tsamou M, Vrijens K, Wang C, Winckelmans E, Neven KY, Madhloum N, de Kok TM, Nawrot TS. Genome-wide microRNA expression analysis in human placenta reveals sex-specific patterns: An ENVIRONAGE birth cohort study. Epigenetics 2020:1–16.

72. Guo S, Huang S, Jiang X, Hu H, Han D, Moreno CS, Fairbrother GL, Hughes DA, Stoneking M, Khaitovich P. Variation of microRNA expression in the human placenta driven by population identity and sex of the newborn. BMC Genomics 2021; 22.

73. Saha B, Ganguly A, Home P, Bhattacharya B, Ray S, Ghosh A, Rumi MAK, Marsh C, French VA, Gunewardena S, Paul S. TEAD4 ensures postimplantation development by promoting trophoblast self-renewal: An implication in early human pregnancy loss. Proceedings of the National Academy of Sciences of the U.S.A. 2020; 117:17864–17875.

74. Fallen S, Baxter D, Wu X, Kim T-K, Shynlova O, Lee MY, Scherler K, Lye S, Hood L, Wang K. Extracellular vesicle RNAs reflect placenta dysfunction and are a biomarker source for preterm labor. Journal of Cellular and Molecular Medicine 2018; 22:2760–2773.

75. Jeon YJ, Choi YS, Rah H, Kim SY, Choi DH, Cha SH, Shin JE, Shim SH, Lee WS, Kim NK. Association study of microRNA polymorphisms with risk of idiopathic recurrent spontaneous abortion in Korean women. Gene 2012; 494:168–173.

76. He Z, Shen F, Qi P, Zha Z, Wang Z. miR-51-3p enhancess the radiosensitivity of prostate cancer cells by inhibiting HSP27 expression and downregulating β-catenin. Cell Death Discovery 2021; 7:1–13.

77. Majem B, Parrilla A, Jiménez C, Suárez-Cabrera L, Barber M, Marín-Guiu X, Alameda F, Romero I, Sánchez JL, Pérez-Benavente A, Rigau M, Gil-Moreno A, et al. MicroRNA-654-5p suppresses ovarian cancer development impacting on MYC, WNT and AKT pathways. Oncogene 2019; 28:6035–6050.

78. Tan Y-W, xu X-Y, Wang J-F, Zhang C-W, Zhang S-C. MiR-654-5p attenuates breast cancer progression by targeting EPSTI1. American Journal of Cancer Research 2016; 6:522–532.

79. Yang L, Wei Q-M, Zhang X-W, Sheng Q, Yan X-T. MiR-376a promotion of proliferation and metastases in ovarian cancer: Potential role as a biomarker. Life Sciences 2017; 173:62–67.

80. Conte FA, Grumbach MM. Disorders of sex determination and differentiation. In: Gardner DG, Shoback DM, Greenspan FS (eds.), Greenspan’s basic & clinical endocrinology. New York: McGraw-Hill Medical; 2007: 479–572.

81. Grumbach MM, Gluckman PD. The human fetal hypothalamus and pituitary gland: the maturation of neuroendocrine mechanisms controlling the secretion of pituitary growth hormone, prolactin, gonadotropins, adrenocorticotropin-related peptides, and thyrotropin. In: Tulchinsky D, Little AB (eds.), Maternal fetal endocrinology, 2nd ed. Philadelphia: Saunders; 1994: 193–262.

82. Muralimanoharan S, Guo C, Myatt L, Maloyan A. Sexual dimorphism in miR-210 expression and mitochondrial dysfunction in the placenta with maternal obesity. International Journal of Obesity 2015; 39:1274–1281.

83. Li H, Ouyang Y, Sadovsky E, Parks WT, Chu T, Sadovsky Y. Unique microRNA Signals in Plasma Exosomes from Pregnancies Complicated by Preeclampsia. Hypertension 2020; 75:762–771.

84. Srinivasan S, Treacy R, Herrero T, Olsen R, Leonardo TR, Zhang X, DeHoff P, To C, Poling LG, Fernando A, Leon-Garcia S, Knepper K, et al. Discovery and Verification of Extracellular miRNA Biomarkers for Non-invasive Prediction of Pre-eclampsia in Asymptomatic Women. Cell Rep Med 2020; 1.

85. Gillet V, Ouellet A, Stepanov Y, Rodosthenous RS, Croft EK, Brennan K, Abdelouahab N, Baccarelli A, Takser L. miRNA Profiles in Extracellular Vesicles From Serum Early in Pregnancies Complicated by Gestational Diabetes Mellitus. J Clin Endocrinol Metab 2019; 104:5157–5169.

86. Sufriyana H, Wu YW, Su EC. Prediction of preeclampsia and intrauterine growth restriction: development of machine learning models on a prospective cohort. JMIR Medical Informatics 2020; 8:e15411.

