## Supplemental File 1 for "Sex differences in microRNA expression in first and third trimester human placenta"

Supplemental File 1: Principal Components Analysis

1A: Fetal Sex Analysis First Trimester PC1 V PC2

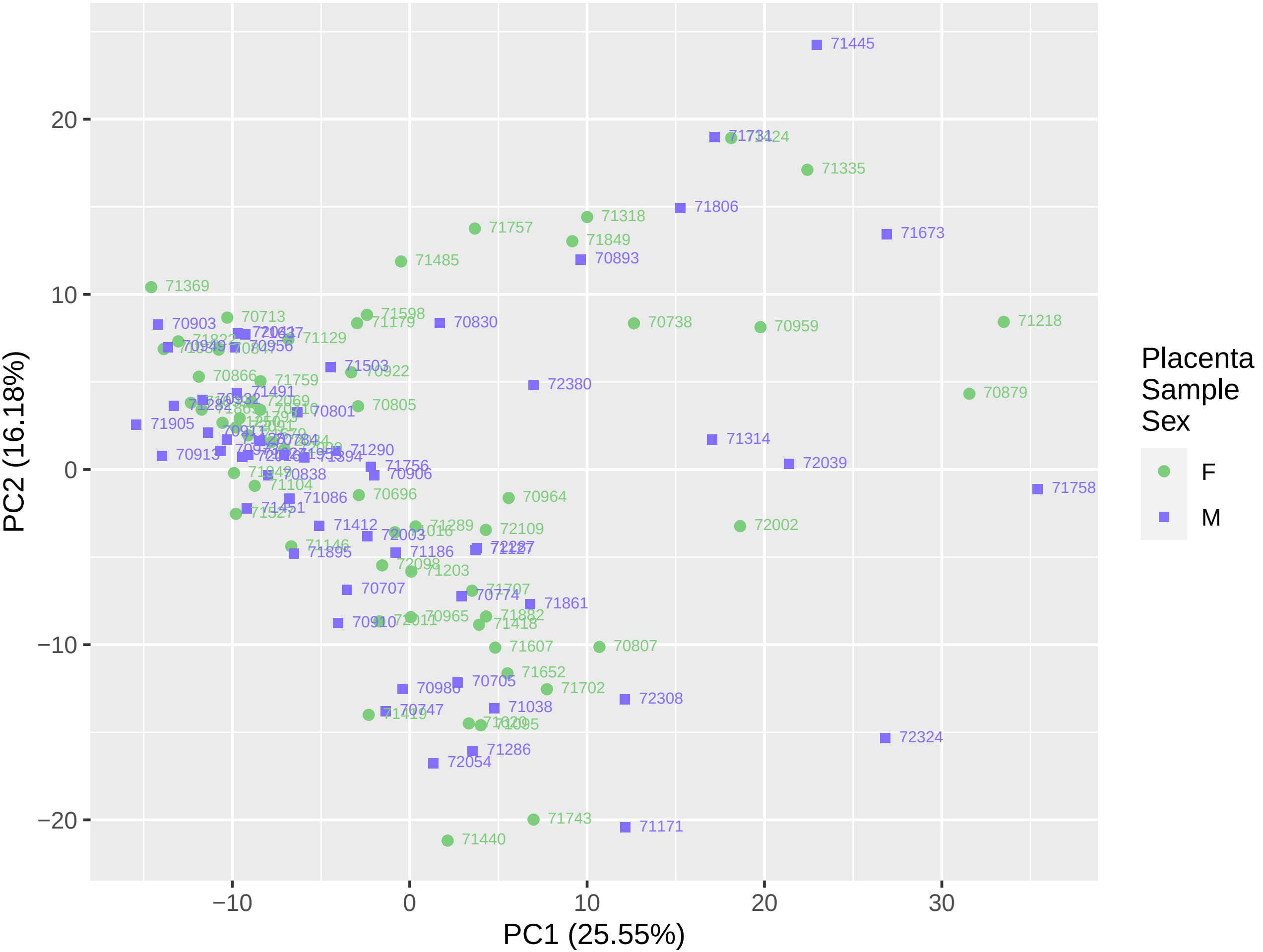

1B: Fetal Sex Analysis First Trimester PC1 V PC3

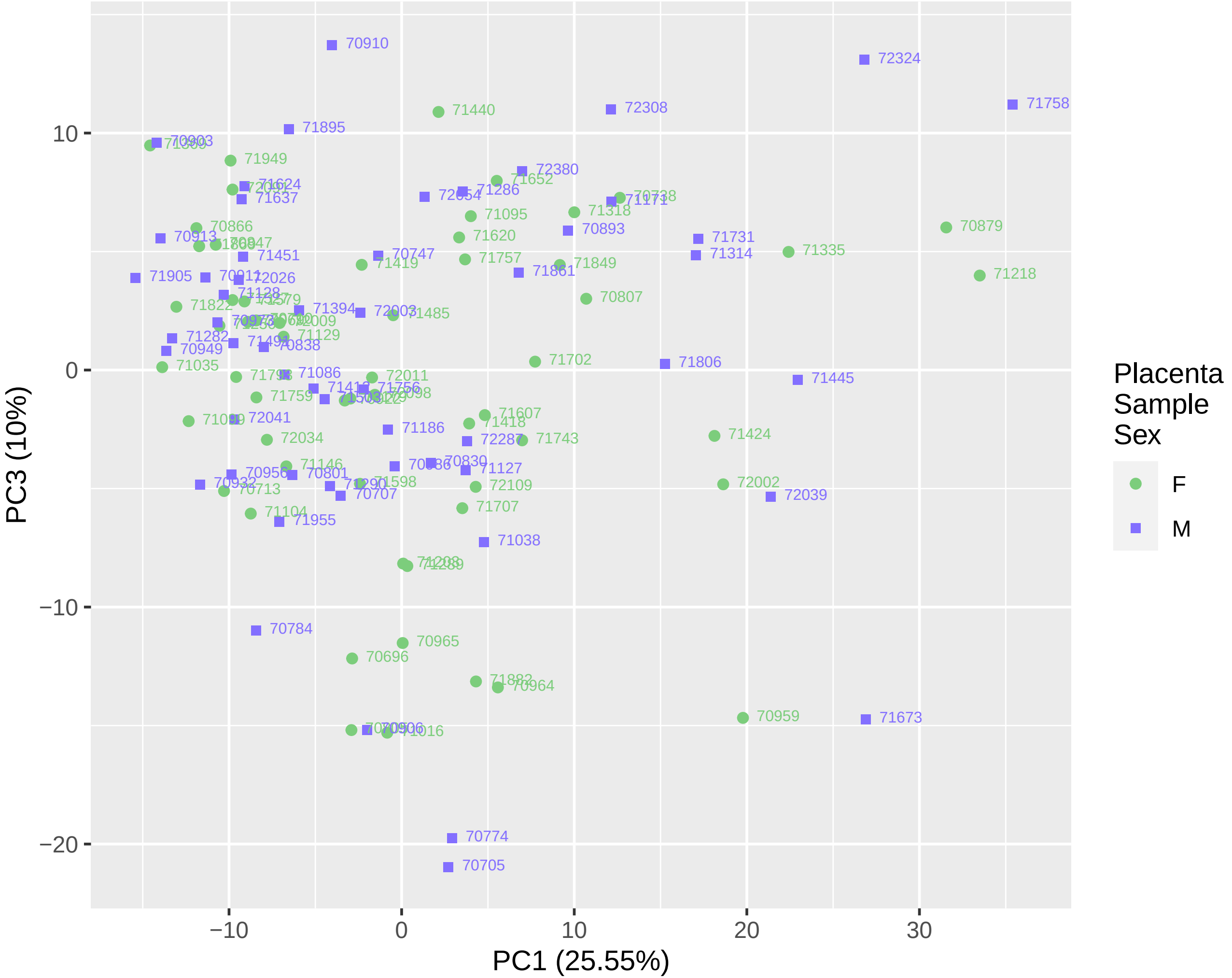

1C: Fetal Sex Analysis First Trimester PC2 V PC3

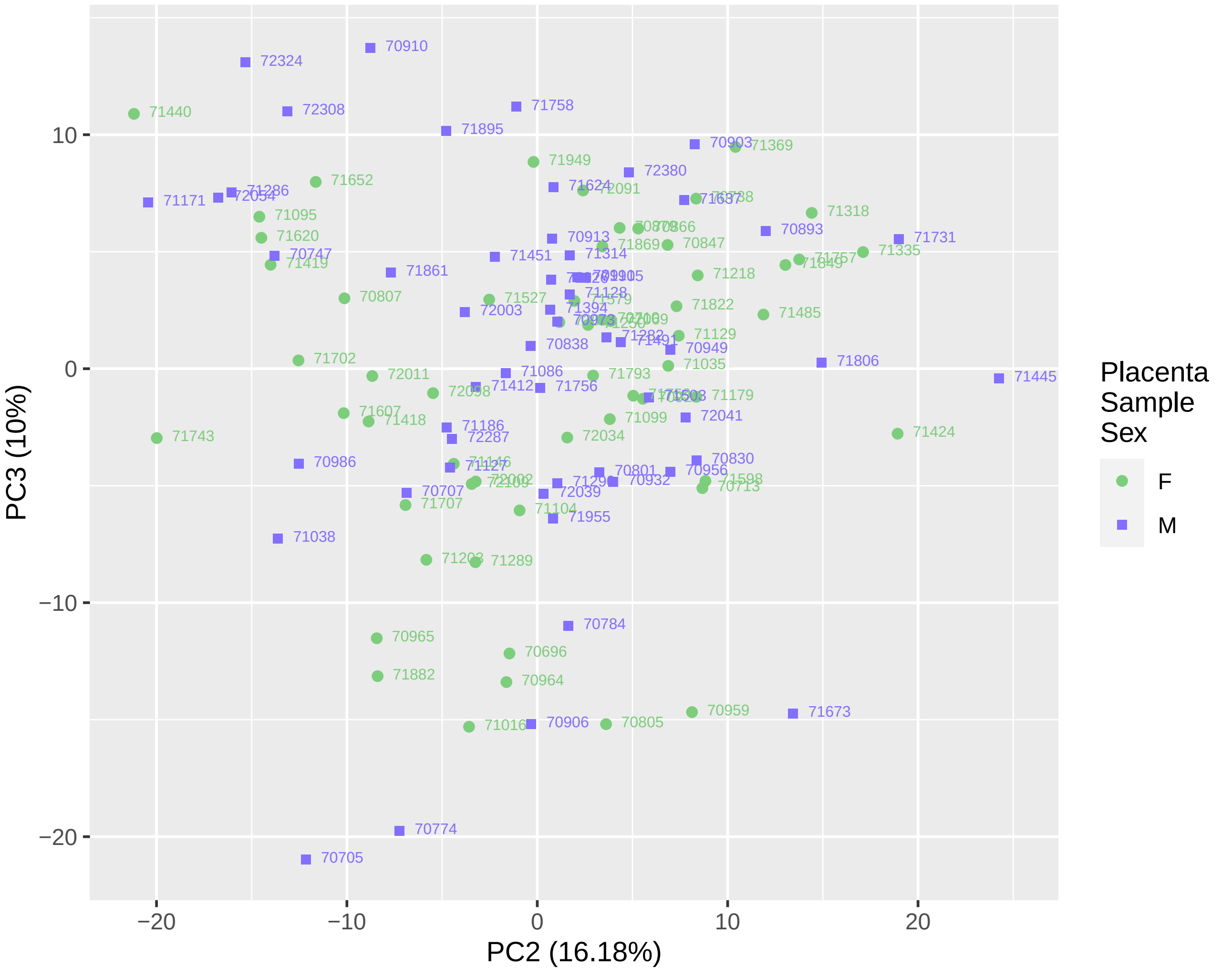

2A: Fetal Sex Analysis Third Trimester PC1 V PC2

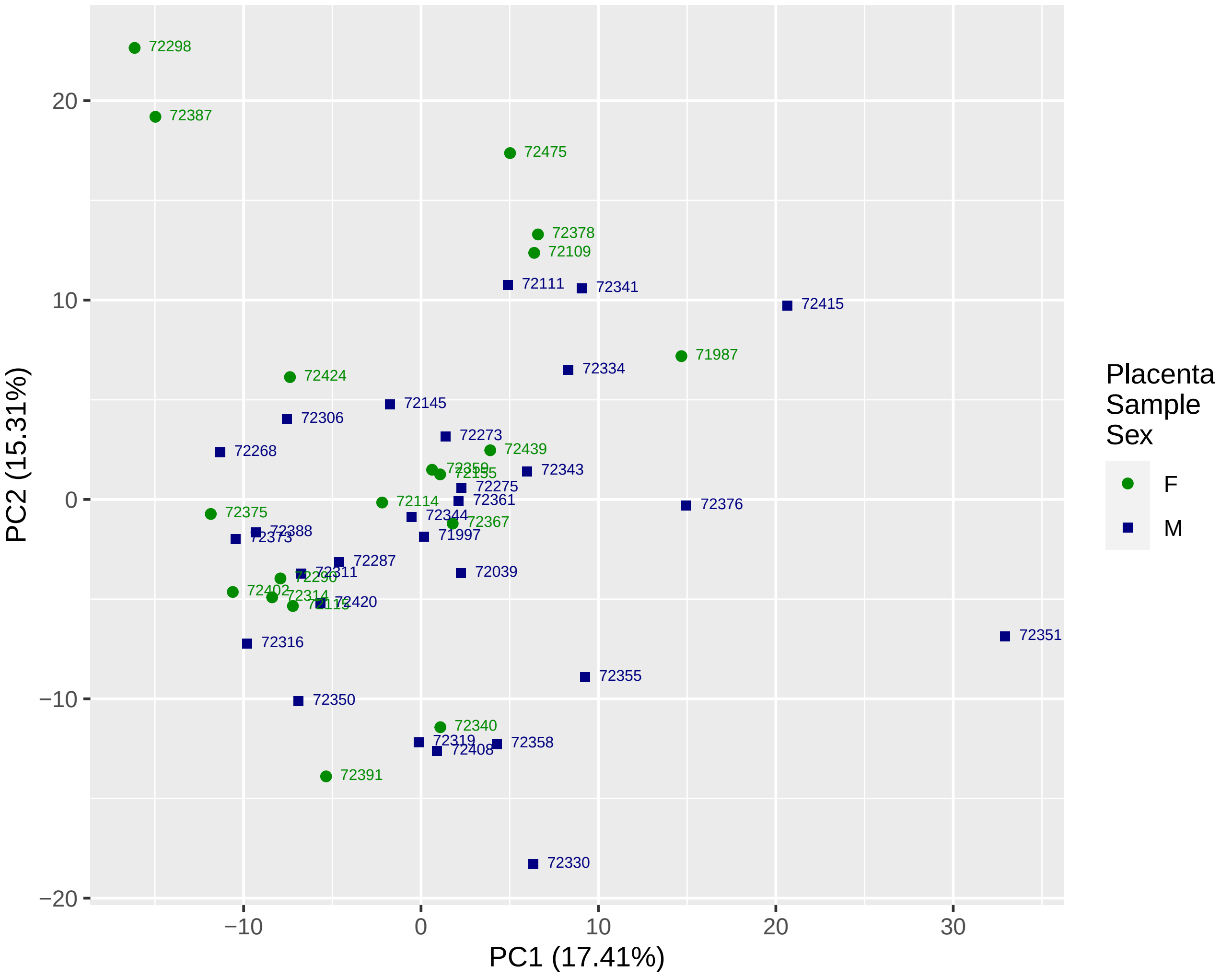

2B: Fetal Sex Analysis Third Trimester PC1 V PC3

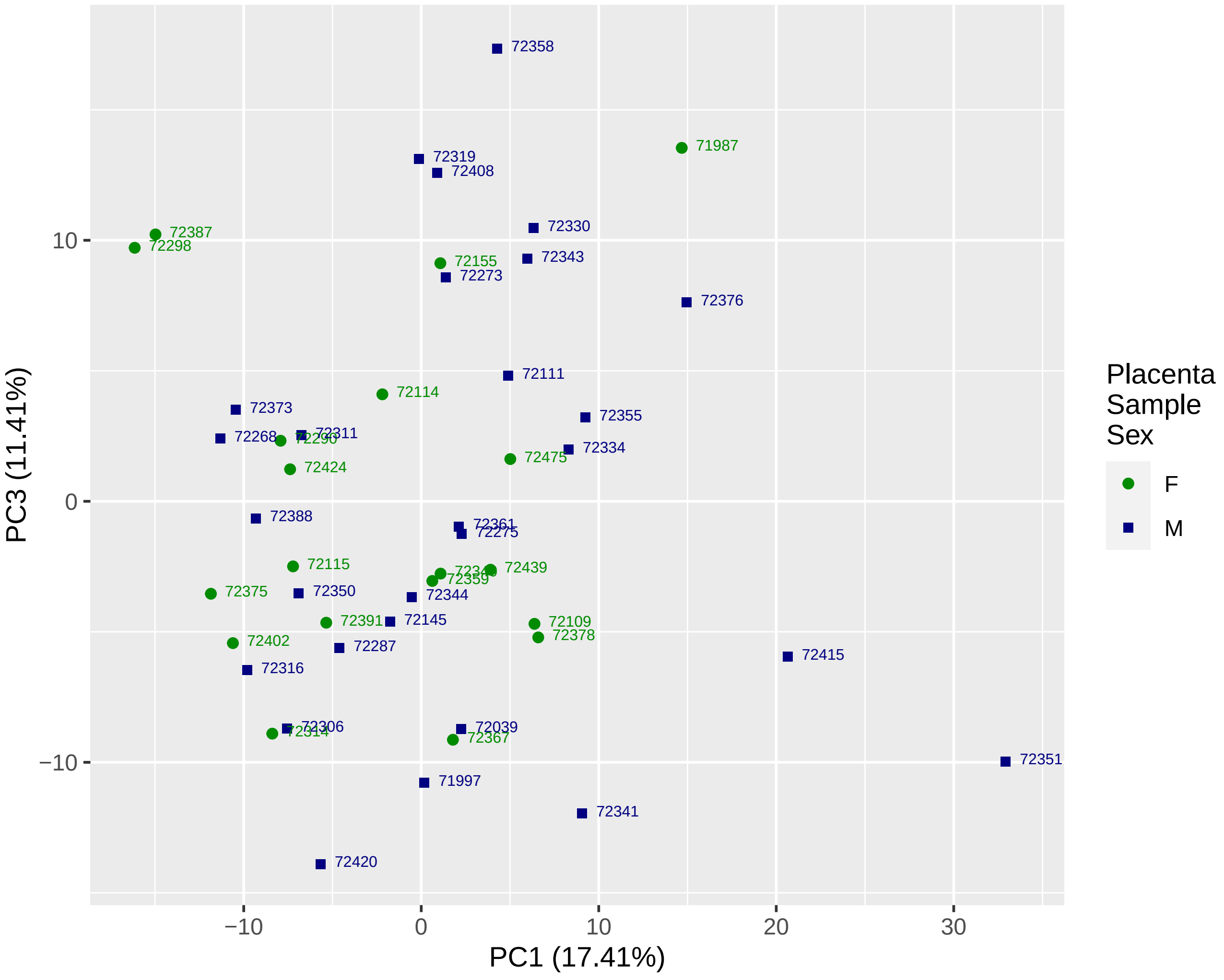

2C: Fetal Sex Analysis Third Trimester PC2 V PC3

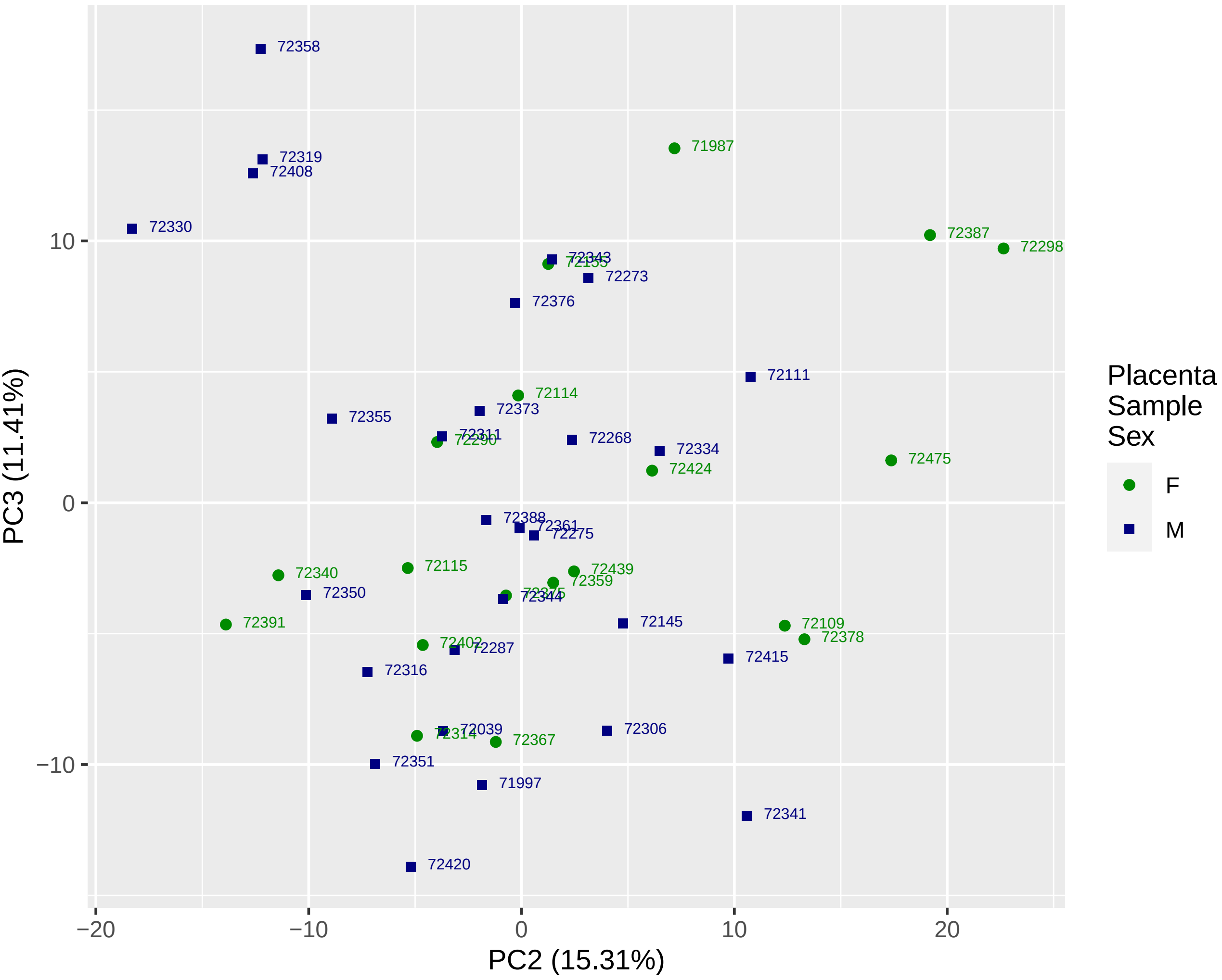

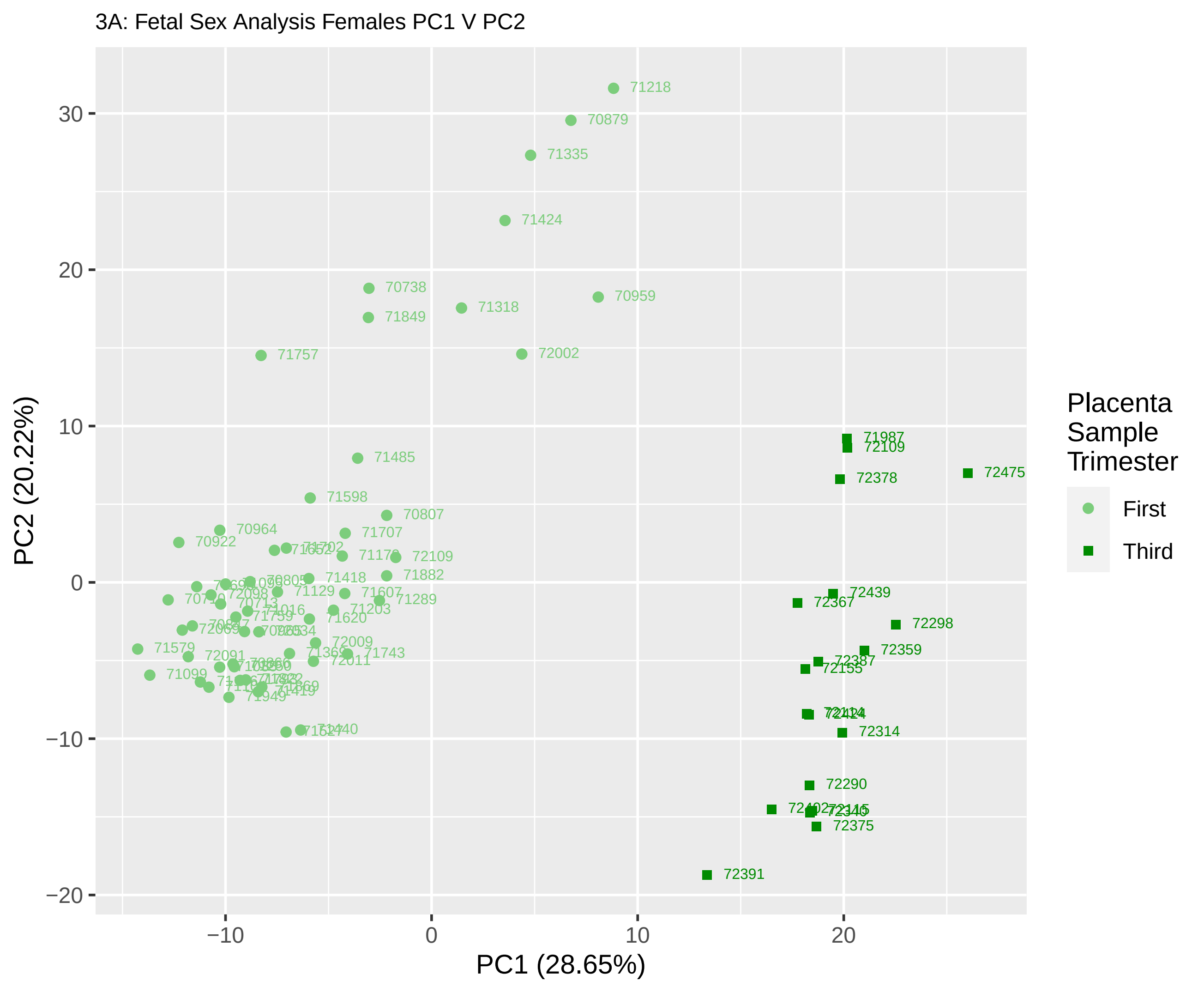
3A: Fetal Sex Analysis Females PC1 V PC2
PC2 (20.22%)
PC1 (28.65%)
30
20
10
0
−10
−20
−10
0
10
20
Placenta Sample Trimester
First
Third
71218
70879
71335
71424
70738
71849
71318
70959
71757
72002
71485
71598
70807
70964
70922
71707
72109
71882
71418
71607
71289
71203
71620
72009
71743
72011
71579
72091
71099
71949
71987
72109
72475
72378
72439
72367
72298
72359
72387
72155
72314
72290
72375
72391

3B: Fetal Sex Analysis Females PC1 V PC3

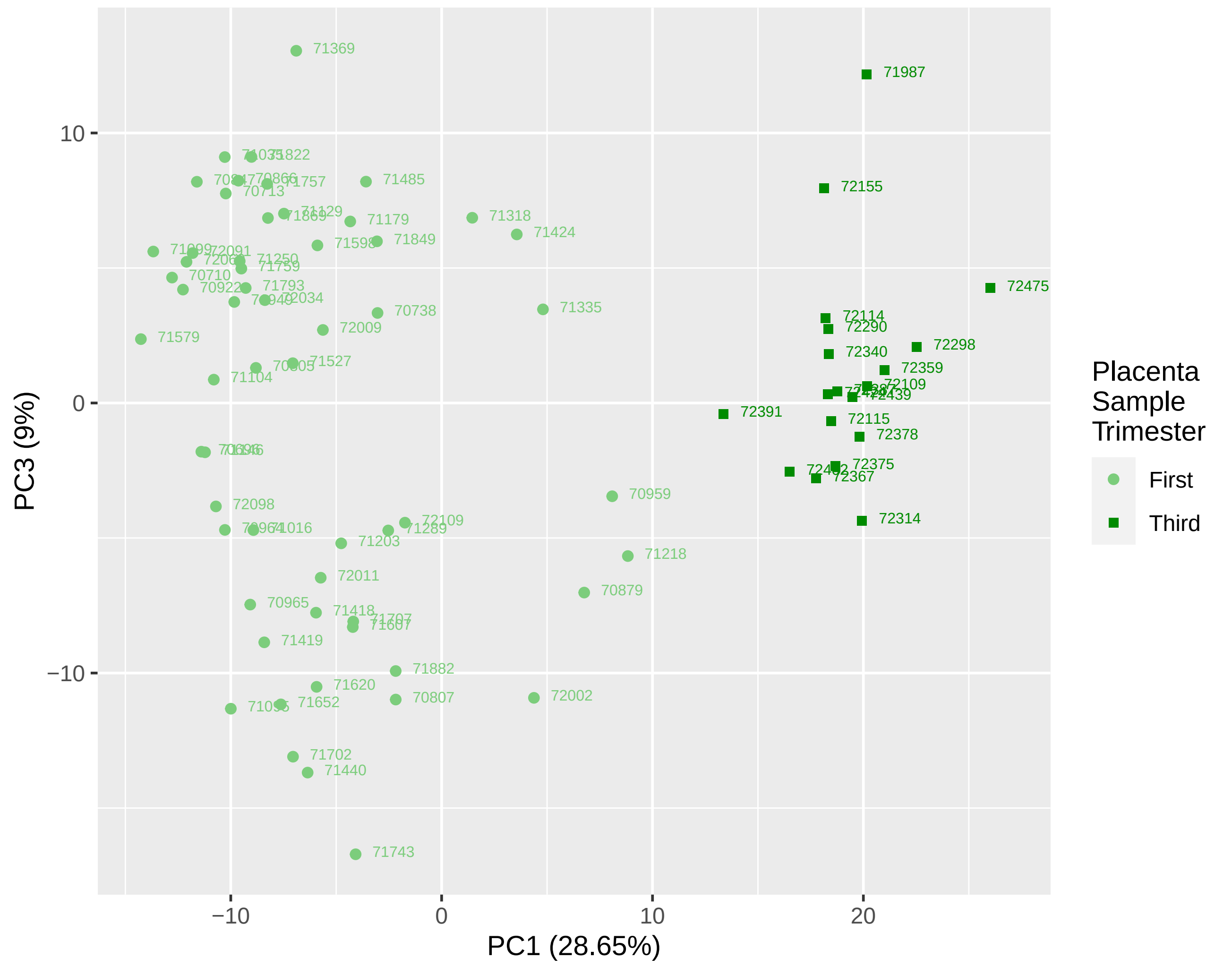

3C: Fetal Sex Analysis Females PC2 V PC3

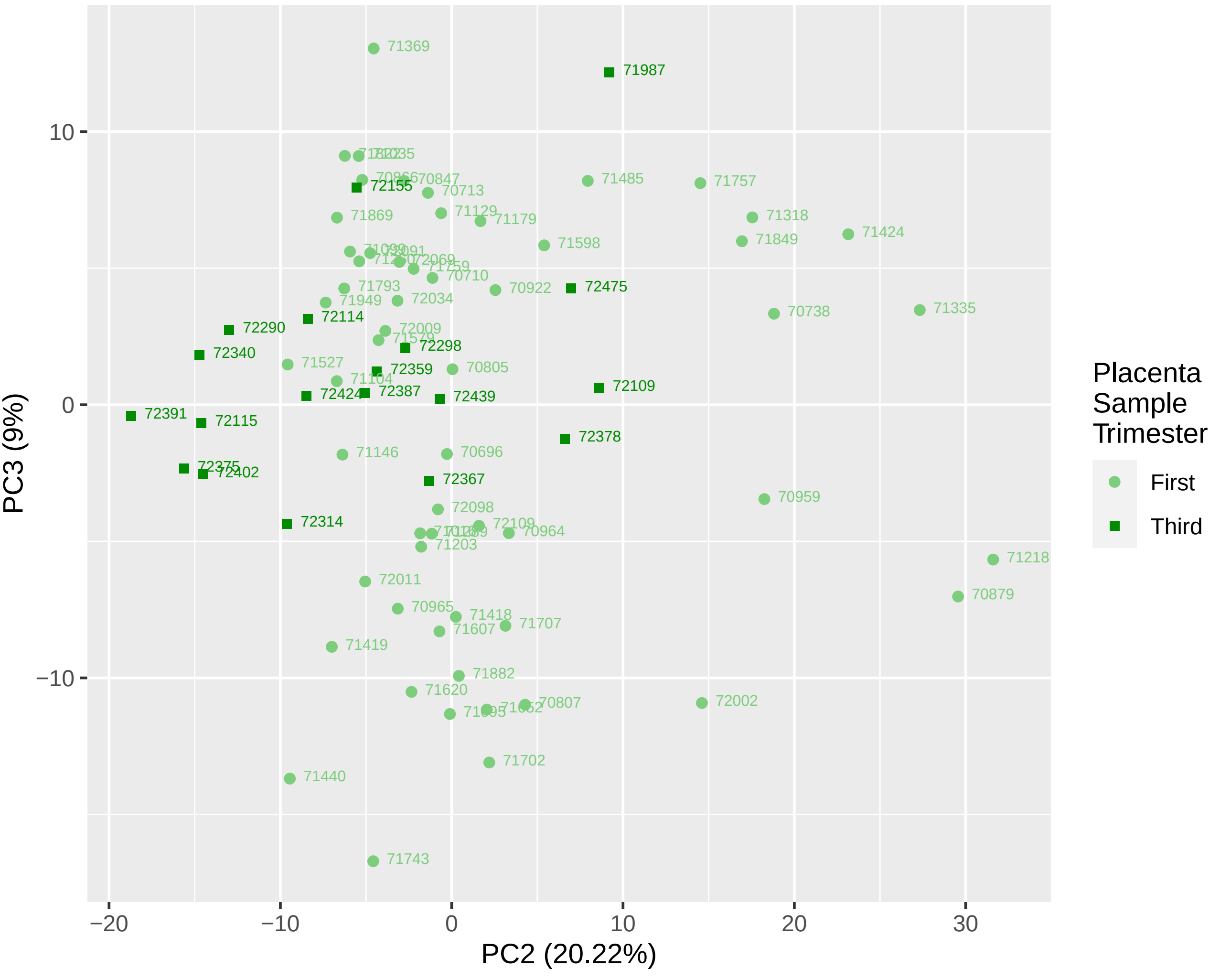

4A: Fetal Sex Analysis Males PC1 V PC2

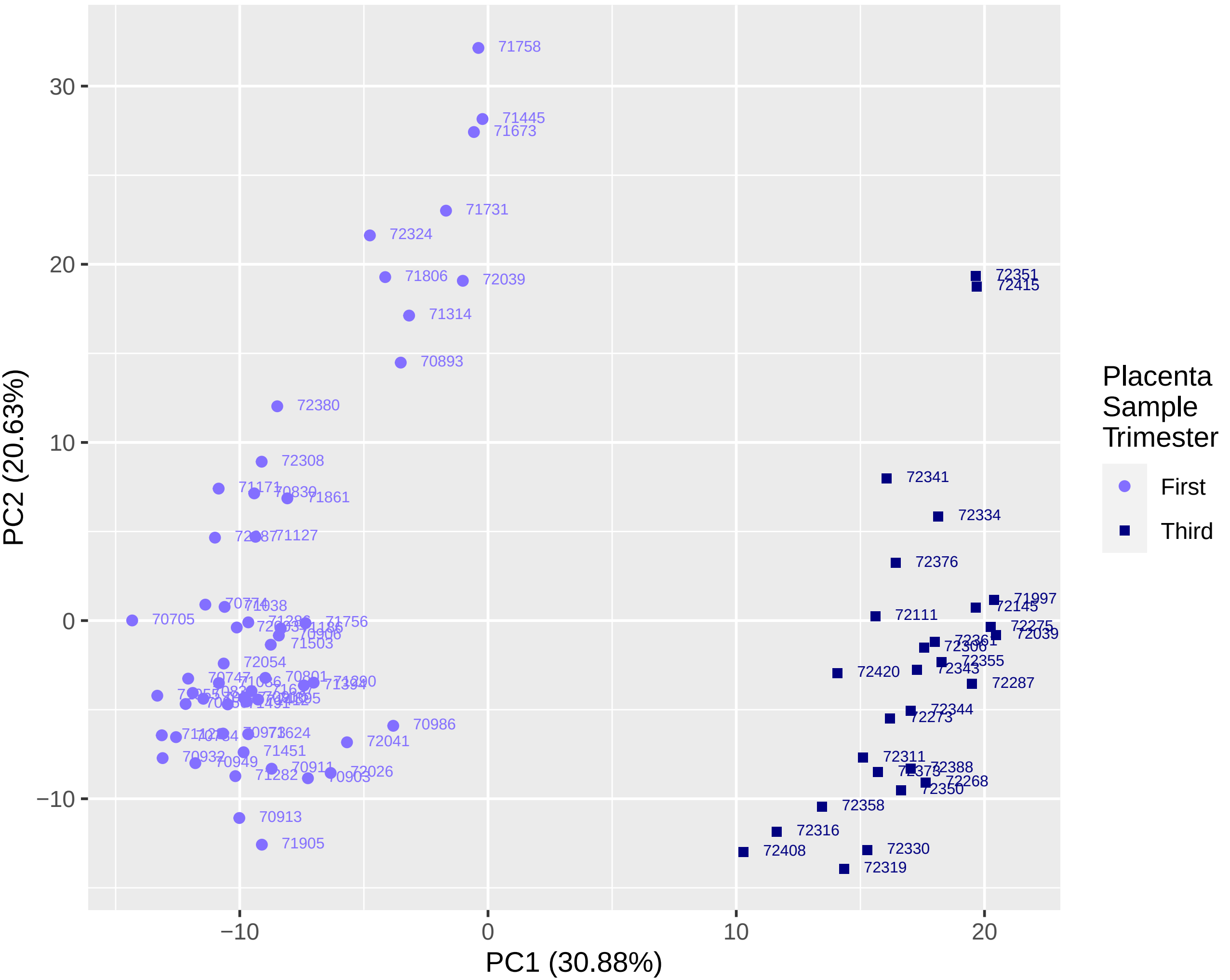

4B: Fetal Sex Analysis Males PC1 V PC3

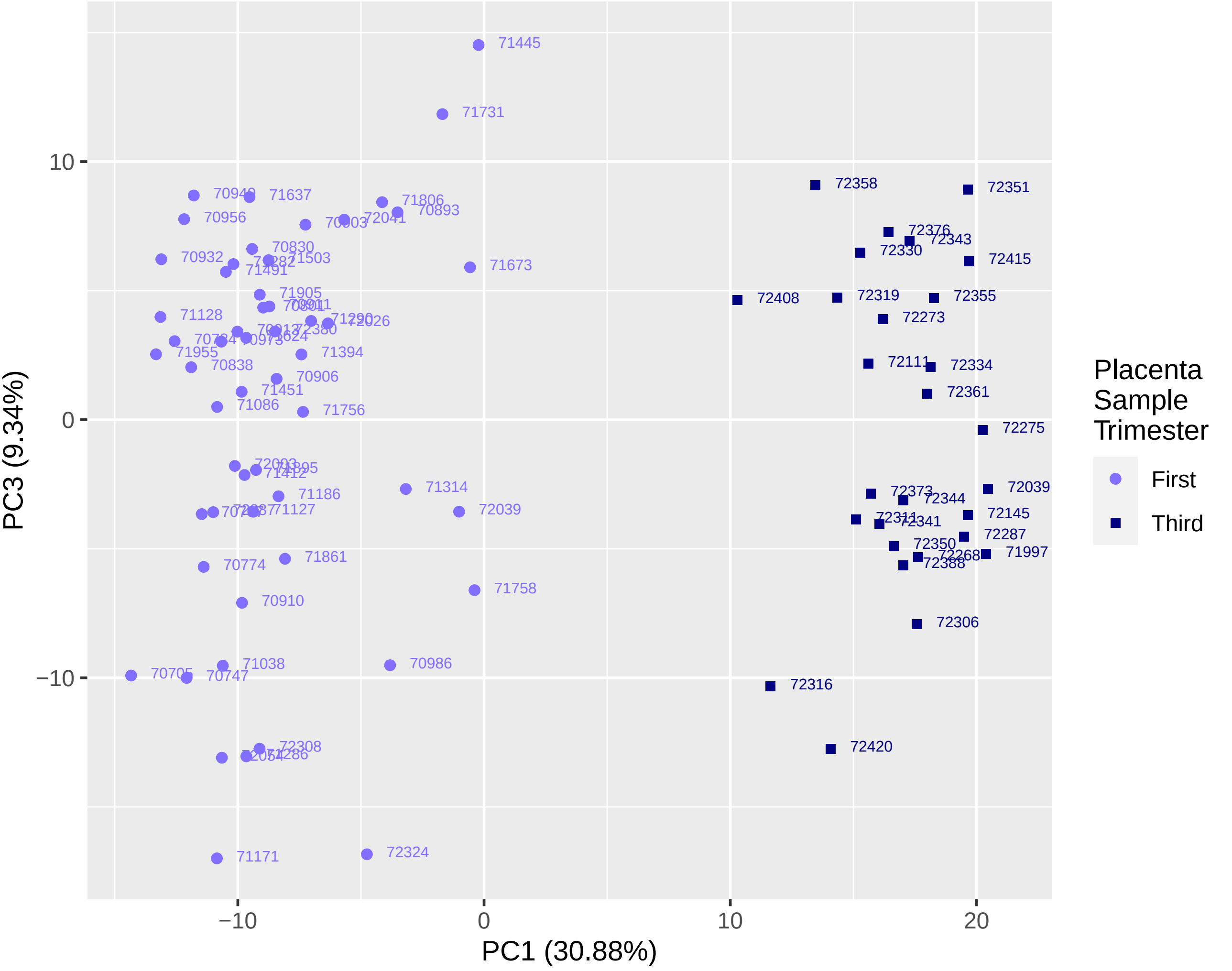

4C: Fetal Sex Analysis Males PC2 V PC3

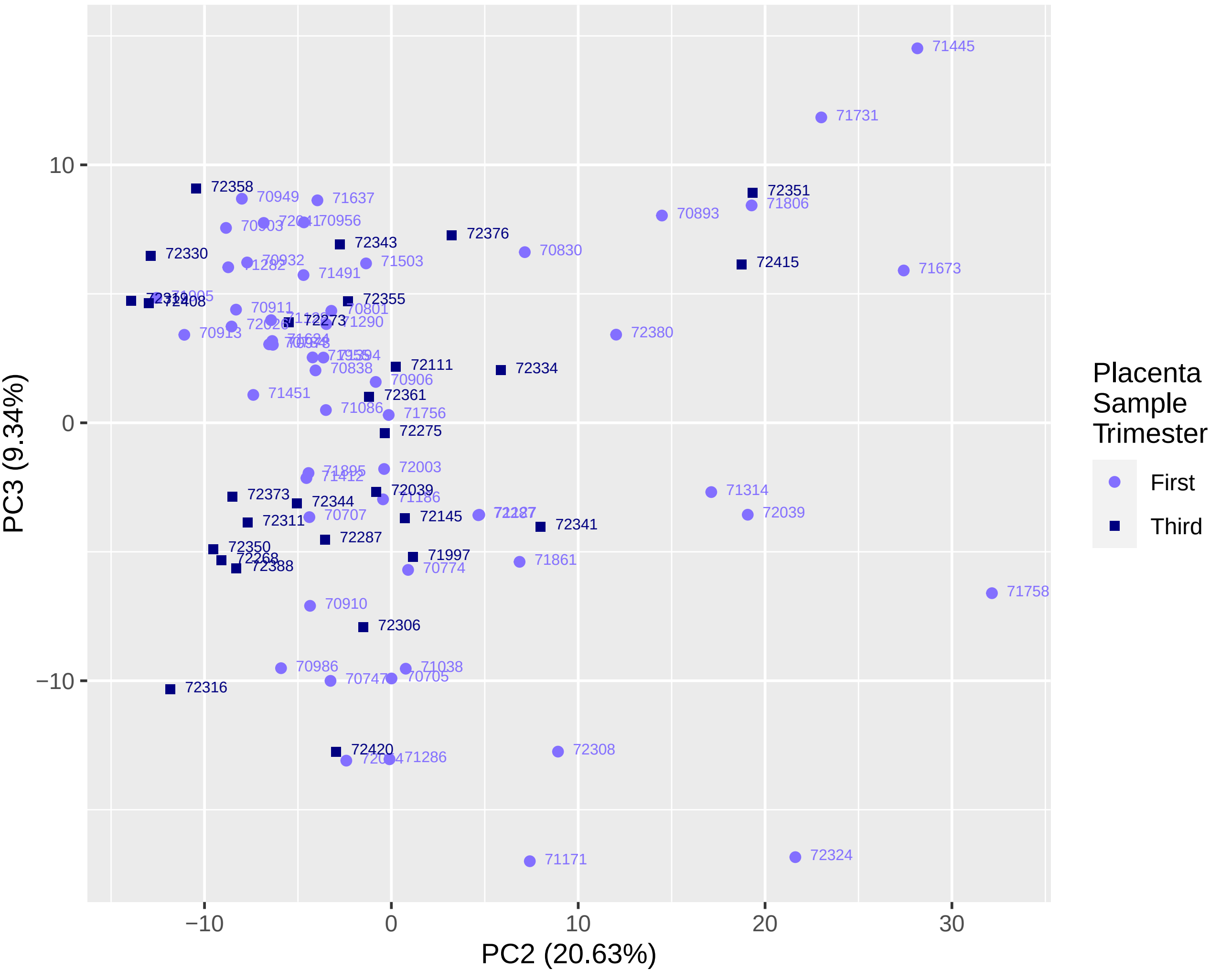
