## Supplementary figures and images for "Sex differences in microRNA expression in first and third trimester human placenta"

### Supplemental File 3

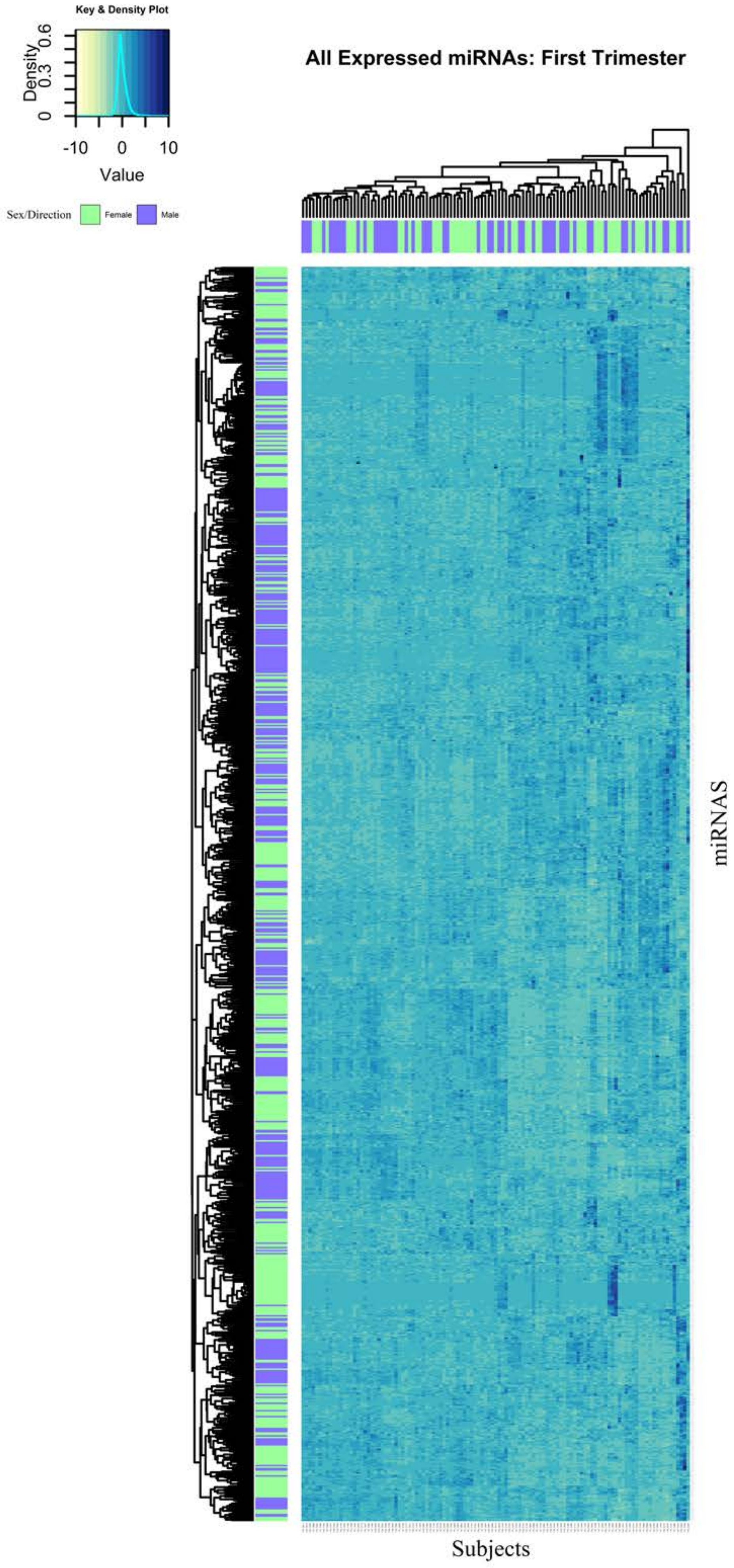

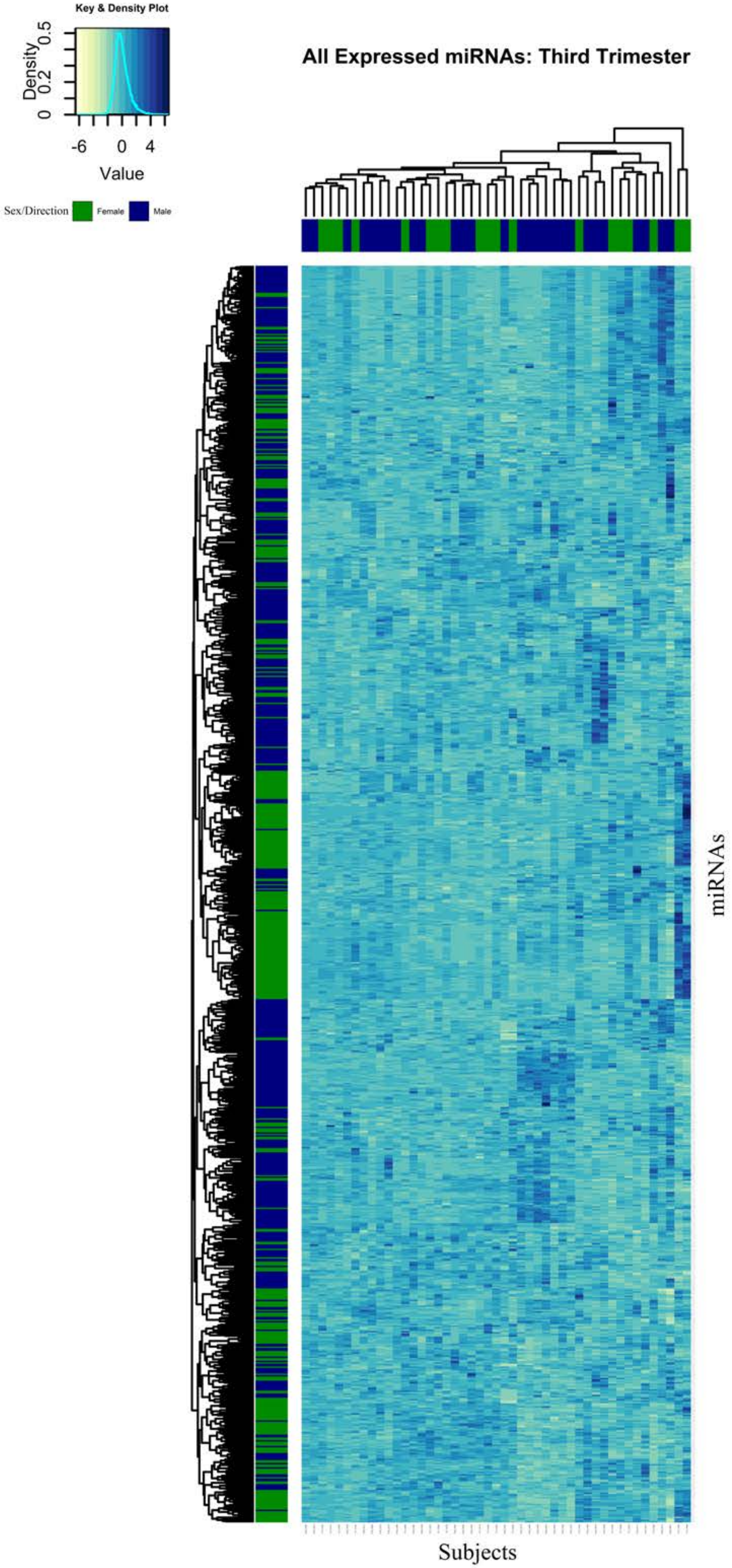

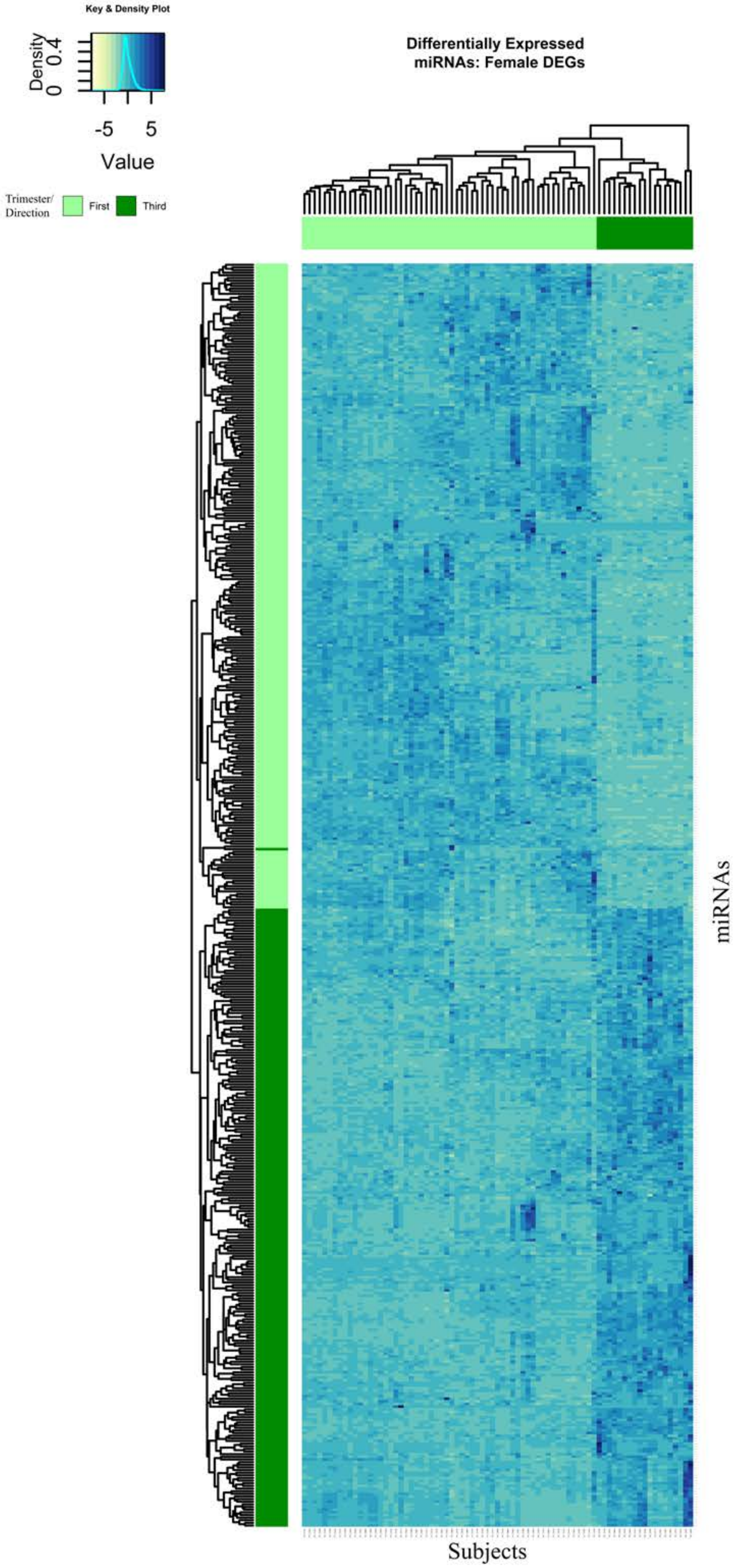

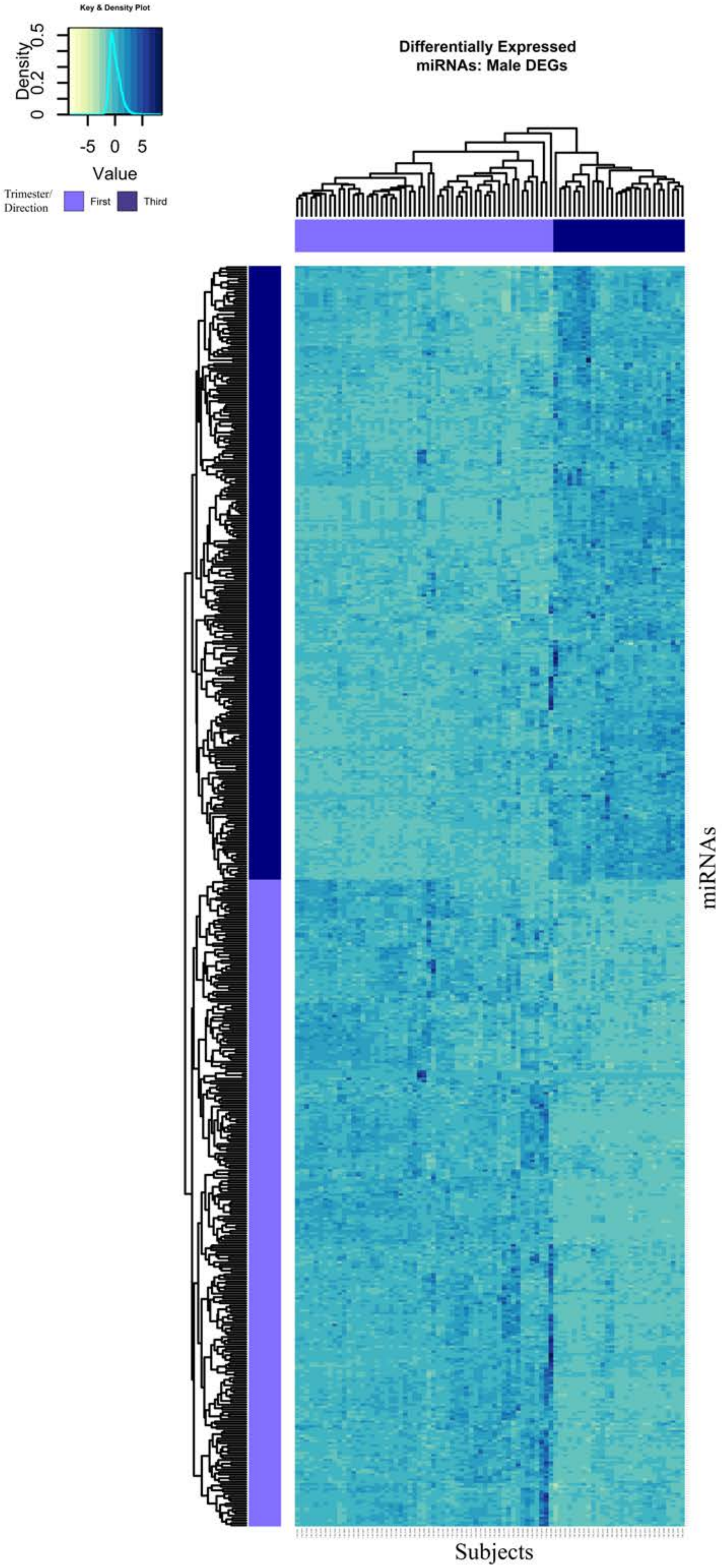
